# Cortico-amygdala synaptic structural abnormalities produced by templated aggregation of α-synuclein

**DOI:** 10.1101/2024.05.15.594419

**Authors:** Nolwazi Z. Gcwensa, Dreson L. Russell, Khaliah Y. Long, Charlotte F. Brzozowski, Xinran Liu, Karen L. Gamble, Rita M. Cowell, Laura A. Volpicelli-Daley

**Affiliations:** Center for Neurodegeneration and Experimental Therapeutics, University of Alabama at Birmingham, Birmingham, Alabama 35294, USA; Department of Psychiatry and Neurobiology, University of Alabama at Birmingham, Birmingham, Alabama 35294, USA; Center for Cellular and Molecular Imaging, Yale University School of Medicine, New Haven, CT 06510, USA

**Keywords:** Parkinson’s disease, Dementia with Lewy Bodies, basolateral amygdala, glutamatergic, presynaptic terminal, synapses, p-α-synuclein

## Abstract

Parkinson’s disease (PD) and Dementia with Lewy bodies (DLB) are characterized by neuronal α-synuclein (α-syn) inclusions termed Lewy Pathology, which are abundant in the amygdala. The basolateral amygdala (BLA), in particular, receives projections from the thalamus and cortex. These projections play a role in cognition and emotional processing, behaviors which are impaired in α-synucleinopathies. To understand if and how pathologic α-syn impacts the BLA requires animal models of α-syn aggregation. Injection of α-synuclein pre-formed fibrils (PFFs) into the striatum induces robust α-synuclein aggregation in excitatory neurons in the BLA that corresponds with reduced contextual fear conditioning. At early time points after aggregate formation, cortico-amygdala excitatory transmission is abolished. The goal of this project was to determine if α-syn inclusions in the BLA induce synaptic degeneration and/or morphological changes. In this study, we used C57BL/6J mice injected bilaterally with PFFs in the dorsal striatum to induce α-syn aggregate formation in the BLA. A method was developed using immunofluorescence and three-dimensional reconstruction to analyze excitatory cortico-amygdala and thalamo-amygdala presynaptic terminals closely juxtaposed to postsynaptic densities. The abundance and morphology of synapses were analyzed at 6- or 12-weeks post-injection of PFFs. α-Syn aggregate formation in the BLA did not cause a significant loss of synapses, but cortico-amygdala and thalamo-amygdala presynaptic terminals and postsynaptic densities with aggregates of α-synuclein show increased volumes, similar to previous findings in human DLB cortex, and in non-human primate models of PD. Transmission electron microscopy showed that PFF-injected mice showed reduced intervesicular distances similar to a recent study showing phospho-serine-129 α-synuclein increases synaptic vesicle clustering. Thus, pathologic α-synuclein causes major alterations to synaptic architecture in the BLA, potentially contributing to behavioral impairment and amygdala dysfunction observed in synucleinopathies.

## Introduction

Although classically thought of as a motor disorder characterized by the clinical presentation of tremor, rigidity, bradykinesia and postural instability, Parkinson’s Disease (PD) has been accepted as a more complex disease with additional presentation of non-motor symptoms including changes in cognition, depression, anxiety, and hallucinations, amongst others (Kumaresan & Khan, 2021; Poewe, 2008). Available treatments only alleviate the motor or non-motor symptoms, but do not address the underlying pathology and progression (Opara et al., 2012; Zhao et al., 2021). One of the major pathological hallmarks of PD includes the presence of Lewy pathology comprised of pathological, insoluble, α-synuclein hyperphosphorylated at serine 129 (p-α-syn). The localization of p-α-syn pathologic aggregates in brain areas such as the cortex, thalamus, and amygdala could contribute to non-motor PD symptoms (Braak & Del Tredici, 2017; Kouli et al., 2018).

The amygdala is an area of the brain that is associated with robust p-α-syn inclusion formation, but has only recently been appreciated for the potential impact it could have on patients’ quality of life (Braak et al., 1994; Carey et al., 2021; Huang et al., 2015; Sorrentino et al., 2019; Yamazaki et al., 2000). The basolateral amygdala (BLA), in particular, is necessary for the association between sensory stimuli and emotional and motivational significance (Tye et al., 2011; W. H. Zhang et al., 2021). The BLA sends projections to the striatum and receives projections from the cortex and thalamus which play a role in cognition and emotional control (Salzman & Fusi, 2010; Šimić et al., 2021). Despite what we know about the presence of α-syn aggregates in the amygdala, relatively little is known about the effects of Lewy pathology on amygdala function. Previous studies have shown that Lewy pathology does not strongly correlate with loss of amygdala volume or cell death in post mortem studies (Harding et al., 2002). However, more recent studies using MRI have shown asymmetrical loss of selective nuclei in the amygdala in PD patients (Kilzheimer et al., 2019; Qu et al., 2024). In addition, hypoconnectivity between regions of the basolateral amygdala (BLA), and related frontal, temporal and insular cortices in MRI studies of PD patients has been reported (Wang et al., 2023). α-Syn aggregation in the amygdala may correlate with changes in amygdala functions.

Animal models of α-syn aggregation can help determine the effect of pathologic α-syn on brain function and behavior. By injecting preformed fibrils (PFFs) into specific brain areas, neurons projecting to the area take up these fibrils which recruit and corrupt endogenous α-synuclein (Luk et al., 2012; Volpicelli-Daley et al., 2011). Bilateral dorsal striatal PFF injections produce robust α-syn aggregate formation in mouse amygdala which associates with reduced performance in fear conditioning thought to be caused at least in part by amygdala dysfunction (Stoyka et al., 2020). This phenotype is not a result of cell death or loss of volume in the mouse amygdala (Stoyka et al., 2020). Electrophysiology studies using the PFF intrastriatal injection model showed defects in the cortico-amygdala, but not thalamo-amygdala excitatory transmission (Chen et al., 2022). These functional and behavioral defects associated with α-syn aggregate formation in mouse amygdala that occur without overt neuron death, suggest other mechanisms of dysfunction associated with p-α-syn aggregation.

The defects in behavior and transmission could be caused by degeneration of synapses in the amygdala caused by α-syn aggregates. A number of neurodegenerative diseases are associated with impairment of synaptic function (Bridi & Hirth, 2018; Taoufik et al., 2018). In PD, loss of nigrostriatal terminals occurs before degeneration of DA neurons (Hornykiewicz, 1998; Kordower et al., 2013). Recent PET imaging studies show synaptic loss in subjects with mild PD symptoms (Matuskey et al., 2020). Additionally, the presence of α-syn micro-aggregate accumulations at the presynaptic terminal correlates with down regulation of presynaptic proteins, such as syntaxin and synaptophysin, as well as postsynaptic density (PSD) proteins, PSD95 and drebrin (Kramer & Schulz-Schaeffer, 2007). Changes in synapses have also been shown in animal models of PD.

Formation of α-syn aggregates corresponds with major loss of dendritic spines in the mouse PFF primary culture model, *in vivo* PFF and in α-syn overexpression mouse models (Blumenstock et al., 2017; Froula et al., 2018). PFF-induced α-syn aggregates also associate with loss of pre-synaptic protein expression and major alterations in molecular signatures of synaptic function (Goralski et al., 2024; Volpicelli-Daley et al., 2011). Thus, formation of α-syn may lead to an early change in the structure of the synapse.

Here, we used the intrastriatal PFF mouse model to determine the effects of α-syn aggregation on synaptic structure in the amygdala. We employed immunofluorescence to label excitatory pre- and postsynaptic puncta in the amygdala. We then developed a method to render three-dimensional surfaces to measure changes in density and volume of synaptic puncta at 6 weeks and 12 weeks following initiation of p-α-syn positive pathological aggregates. We show that the density of excitatory cortico-amygdala or thalamo-amygdala synapses is not altered at either time point. However, pre- and postsynaptic puncta that contain p-α-syn show increased volume in both cortico- and thalamic-amygdala synapses.

## Materials and Methods

Unless otherwise noted, all materials were purchased from Fisher Scientific.

### Animals

The Institutional Animal Care and Use Committee at the University of Alabama at Birmingham approved all animal protocols IACUC 22112 (06/22/2022 – 06/21/2023), IACUC 22447 (02/27/2023 – 01/06/2025) and IAUCUC 22614 (10/18/2022 – 10/18/2023). C57BL/6J mice (Strain 000664) were purchased from Jackson Labs and maintained on a 12-hour light/dark cycle with unrestricted access to food and water. Both male and female mice were included in each study, unless otherwise stated.

### Preparation of recombinant α-synuclein PFFs

Mouse monomeric α-synuclein was purified in E. Coli and a Pierce LAL high-capacity endotoxin removal resin was used to minimize endotoxin as previously described (Volpicelli-Daley et al., 2014). PFFs (5mg/mL) were generated from monomeric α-synuclein through agitation in 150 mM KCl/50 mM Tris-HCl at 37 °C for 7 days (Bousset et al., 2013; Stoyka et al., 2020). PFFs were stored at −80°C until use. Immediately before injection, PFFs were sonicated using a Q700 Sonicator with circulating water at 15 °C. Samples at 22 µL were placed in a 1.5 mL sonicator tube (Active Motif, NC0869649) and sonicated for 15 min (amplitude 30; pulse on 3 s; pulse off 2 s). Fragmentation of PFFs between 20 and 70 nm fragments was confirmed using dynamic light scattering.

### Intrastriatal injection of recombinant α-syn fibrils

Three- to four-month old mice were placed on a stereotactic frame under deep anesthesia with vaporized isoflurane. Thereafter, mice were bilaterally injected with 2 μL (5mg/mL) per side of either sonicated fibrils (10 µg total protein), monomeric α-synuclein (10 µg total protein) or phosphate-buffered saline (PBS). The injections were carried out at a rate of 0.5 μL/min after which the needle remained in place for 5 minutes before a gradual withdrawal. To target the dorsal striatum, mice were injected at coordinates +1.0 mm anterior-posterior, ±2.0 mm mediolateral, and −3.2 mm dorsoventral measured from dura.

### Immunofluorescence and immunohistochemistry

At 6- or 12-weeks post-injection, mice were anesthetized with isoflurane and underwent transcardial perfusion with a 0.9% saline solution containing 10 units/mL heparin and sodium nitroprusside (0.5% w/v) followed by a cold 4% paraformaldehyde (PFA) solution in PBS. Thereafter, brains were harvested and postfixed in the same 4% PFA in PBS solution for 12 hours at 4 °C then submerged in cryoprotectant (30% sucrose in PBS) for 24–48 hours and snap frozen in a dry ice/ethanol slurry for storage at −80 °C. Brains were sectioned at 40 μm thickness on a freezing microtome (Leica SM 2010 R). Serial sections of the brains were placed in a 6-well to ensure each well represented the entire forebrain with slices spaced 240 μm apart. Brain sections were stored in a solution of 50% glycerol, 0.01% sodium azide in tris-buffered saline (TBS).

For immunofluorescence, sections were rinsed three times in cold TBS (5 min) then incubated in a solution of 10 mM sodium citrate, 0.05% Tween-20 (pH 6.0) for 1 h at 37 °C for antigen retrieval. Thereafter, sections were rinsed three times with cold TBS (5 min) then blocked and permeabilized in a solution of 5% normal goat serum, 0.01% TritonX-100 in TBS for 1 h at 4 °C with agitation. Sections were then incubated in a solution of 5% normal goat serum in TBS with primary antibodies (Table 1) for 16 h at 4 °C with agitation. Thereafter, sections were rinsed as before (3 x 5 min, cold TBS) then incubated with AlexaFluor conjugated secondary antibodies (Thermofisher) in 5% normal goat serum in TBS for 2 h at 4°C with agitation. Brains were again rinsed with cold TBS (3 x 5 min). To stain nuclei, Hoechst was included in the second TBS wash (1:500). Finally, sections were incubated in solution of cupric sulfate (10 mM) and ammonium acetate (50 mM) for 10 minutes at room temperature with agitation to quench autofluorescence from lipofuscin before quenching the reaction with two 5 min rinses in distilled water. Finally, sections were stored in TBS before mounting using Prolong Gold (Thermofisher).

**Table 1:** Primary Antibodies.

| ANTIBODY TARGET | Supplier | Catalog No. | RRID | Host species/Isotype/Mono or Polyclonal | Dilution |
| --- | --- | --- | --- | --- | --- |
| **VGLUT1** | Synaptic Systems | 135 304 | AB_887878 | Guinea Pig/IgG/Poly | 1 : 1 000 |
| **VGLUT2** | Synaptic Systems | 135 421 | AB_2619823 | Mouse/IgG/Mono | 1 : 1 000 |
| **HOMER1** | Synaptic Systems | 160 006 | AB_2631222 | Chicken/IgG/Poly | 1 : 1 000 |
| **PSER129-SYNUCLEIN** | Abcam | 51253 | AB_869973 | Rabbit/IgG/Mono | 1 : 5 000 |

### Wide Field Fluorescence Microscopy

For visualization of pS129-positive aggregates in whole hemispheres containing the amygdala and the thalamus, sections were imaged as tiled files using an inverted Zeiss Axiovert.Z1 microscope. Carl Zeiss software was used to stitch single images acquired with LD Plan-Neofluar 20X/0.4 Corr M27 air objective. Single frame images of amygdala and thalamus were acquired using the LD Plan-Neofluar 20X/0.4 Corr M27 air objective and LD Plan-Neofluar 20X/0.6 Corr air objective.

### Confocal Microscopy and Imaris 3D surface reconstruction

#### Image acquisition

Imaging in the BLA was performed on coronal sections corresponding to levels −1.22 mm and −1.94 mm relative to bregma. Hoechst nuclear stain of the external and amygdala capsules was used to identify the BLA. Images were collected using a Nikon Ti2 confocal microscope using a Nikon CFI Plan Apochromat λD 60X/1.42 oil objective with laser power, gain, offset, resolution, and pinhole diameter, kept consistent across all mice for each experiment. Eight to ten z-stack frames (step size 0.125 µm, steps > 20) were imaged from two sections for each mouse. *Image processing:* Images were deconvolved using the Richardson-Lucy algorithm at 40-iterations using NIS Elements Imaging Software. *Generate 3D Surfaces:* For each pre- and post-synaptic element, a specific layer was created using Imaris Software with parameters optimized as outlined in Table 2. To assess whether inclusion formation had an effect of the number of pre- and post-synaptic puncta that are closely juxtaposed, presynaptic puncta with centers that were within 0.01 μm distance from at least one postsynaptic puncta center were filtered. Synaptic loci were defined according to the distance between the centers of a presynaptic surface A and postsynaptic surface B, such that distance from center A to center B < 0.01 µm = synaptic loci. Any pre- and postsynaptic surfaces that did not fit this definition were defined as non-synaptic loci. To quantify the density of synaptic loci, the number of synaptic puncta in a population were counted and normalized to the volume of the frame (29,175 µm for all VGLUT1 experiments and 34,410 µm for all VGLUT2 experiments). To determine the number of synaptic puncta that contain p-α-syn inclusions, a 3D surface was generated for p-α-syn according to surface parameters indicated in Table 2. Thereafter, synaptic surfaces were defined as positive for p-α-syn if the distance between the center of synaptic surface A, and the p-α-syn surface center B < 0.1 µm. Synaptic surfaces that did not fit into this definition were considered negative for p-α-syn. Thereafter, we compared the mean volumes for synaptic puncta populations that contained p-α-syn (w p-α-syn) and those that did not contain p-α-syn (wo p-α-syn).

**Table 2:** 3D surface parameters.

| **SURFACE** | Surface Grain Size (μm) | Diameter of Largest Sphere (μm) | Threshold Value (% of Total Signal) | Filter Surfaces: Voxels | Filter Surface: Volume |
| --- | --- | --- | --- | --- | --- |
| **VGLUT1** | 0.05 | 0.70 | 90 | > 10 | < 2.00 μm<sup>3</sup> |
| **VGLUT2** | 0.20 | 0.40 | 90 | > 10 |  |
| **HOMER1** | 0.15 | 0.30 | 90 | > 10 | < 2.00 μm<sup>3</sup> |
| **PSER129-SYNUCLEIN** | 0.05 | 0.60 | 95 |  |  |

### Transmission electron microscopy

Male mice (n = 4 PFF, n = 4 PBS) were anesthetized with isoflurane and perfused with PBS followed by 2.5% glutaraldehyde and 2% paraformaldehyde in PBS (pH 7.5) at room temperature using a peristaltic pump at 3 mL/min). Brains were removed and immersion fixed in 2.5% glutaraldehyde and 2.0% paraformaldehyde in PBS for 2 hours at room before overnight storage at 4°C overnight. Brains were embedded in 2% 255-bloom calf skin gelatin with 3% agarose in room temperature PBS. Sections were cut on a vibratome at 200 µm thickness in room temperature 1XPBS. The BLA was dissected and placed in a 1.5 mL tube with 2% paraformaldehyde in 0.1 M cacodylate buffer at pH 7.4 and stored at 4 °C. The samples were shipped in 0.1 M phosphate buffer to Center for Cellular and Molecular Imaging EM Core facility at Yale Medical School.

Mouse brain tissue was further post-fixed in 1% OsO4 and 0.8% potassium ferricyanide in 0.1 M cacodylate buffer at room temperature for one hour. Specimens were then en bloc stained with 2% aqueous uranyl acetate for 30 minutes, dehydrated in a graded series of ethanol up to 100%, substituted with propylene oxide, and embedded in EMbed 812 resin (Electron Microscopy Sciences, Hartfield, PA). Sample blocks were polymerized in an oven at 60°C overnight. Ultrathin sections (60 nm) were cut using a Leica ultramicrotome (UC7) and post-stained with 2% uranyl acetate and lead citrate. The sections were examined with an FEI Tecnai transmission electron microscope at an 80 kV accelerating voltage, and digital images were acquired with an AMT NanoSprint15 MK2 camera (Advanced Microscopy Techniques, Woburn, MA).

To analyze TEM images, excitatory, asymmetrical synapses were identified. The following exclusion criteria were applied: poorly defined or unclear PSDs, SVs or pre- and postsynaptic membranes; presynaptic compartment with fewer than 4 SVs; pre- and postsynaptic compartments at the edge of the frame of collection cutting through SVs/PSDs; synapses where convolution neural network algorithm applied via Python Software identified < 85 % of manually counted SVs. *To measure PSD lengths and SVs counts*: PSD was manually traced and measured using the segmented line tool in ImageJ Software. SVs vesicles were counted manually. *To count the number of docked SVs per PSD length:* A line was manually traced of the PSD juxtaposed to presynaptic compartment of interest. Docked vesicles were defined as SVs the fell within distance ≤ 100 nm from active zone adjacent to traced PSD length and counted manually. *To measure the area of synaptic vesicles and intervesicular distance to nearest neighbor:* Presynaptic compartments of interest were cropped and analyzed using convolution neural network algorithm trained on mouse synapses operated via Python Software (Imbrosci et al., 2022). All analyses were performed in a blinded manner.

### Statistical and Graphical Analysis

Analyses and graphs were generated using SPSS or GraphPad Prism. The mean density of synaptic puncta for each frame per animal was averaged to generate one mean density value for each mouse. To compare the means between three groups, PBS, monomer or PFF, ordinary one-way ANOVA was used. Normality between groups was assessed using the Shapiro-Wilkes measure and standard variance was determined using the Browne-Forsythe Test to assess the equality of variances in ANOVA analyses. If one-way ANOVA revealed a statistical difference between means of at least two of the three groups, Tukey’s multiple comparison test with single pooled variance was calculated to determine which specific groups were statistically different. Where the variance between groups was not equal, the Brown-Forsythe correction was applied. For two group-analyses, means were compared using independent student’s t-test and normality between groups was assessed using the Shapiro-Wilkes measure. F-test was employed to assess the equality of variances between two groups and where variances were not equal, the Welch’s t-test was employed. For electron microscopy, values that were more than 2 standard deviations from the mean were excluded. Analyses of the PSD length and docked vesicles divided by the PSD length revealed right-skewed data. The data were thus transformed using Log10. Data were analyzed using linear mixed model with mouse number as “subjects” and synapse number as repeated measures, compound symmetry structure was used with treatment as a fixed effect. For the synaptic vesicle area, The “AI” algorithm for analyses of synaptic vesicle area thresholded the data such that the data were not continuous (Imbrosci et al., 2022). Therefore, a Fisher’s exact test was performed comparing the percentage of cases that were above or below the grand median for PBS or PFF.

## Results

### PFF injection into the striatum induces robust α-synuclein inclusion formation in mouse BLA at 6- and 12-weeks post-injection

To examine the extent and localization of aggregation at 6- and 12-weeks post-injection an antibody which recognizes α-syn phosphorylated at Ser129, was used to identify α-syn inclusions. Mice injected with monomeric α-syn or phosphate buffered saline (PBS) were used as negative controls as these injections do not induce α-syn inclusion formation (Supplemental Figure 1). At 6- and 12-weeks following intrastriatal PFF injection there was robust formation of phosphorylated α-syn aggregates in the basolateral amygdala (BLA (Figure 1 A, D). α-Syn aggregates also appeared in cortical layer 5 and the paraventricular nucleus of the thalamus at 6- and 12-weeks post-injection (Figure 1A, D and Supplemental Figure 2). Double labelling with phosphorylated α-synuclein marker pSer129-α-syn and Hoechst nuclear marker showed somal and neuritc inclusions as well as small aggregates at low and high magnifications in the BLA (Figure 1B, C and E, F).

**Figure 1.**
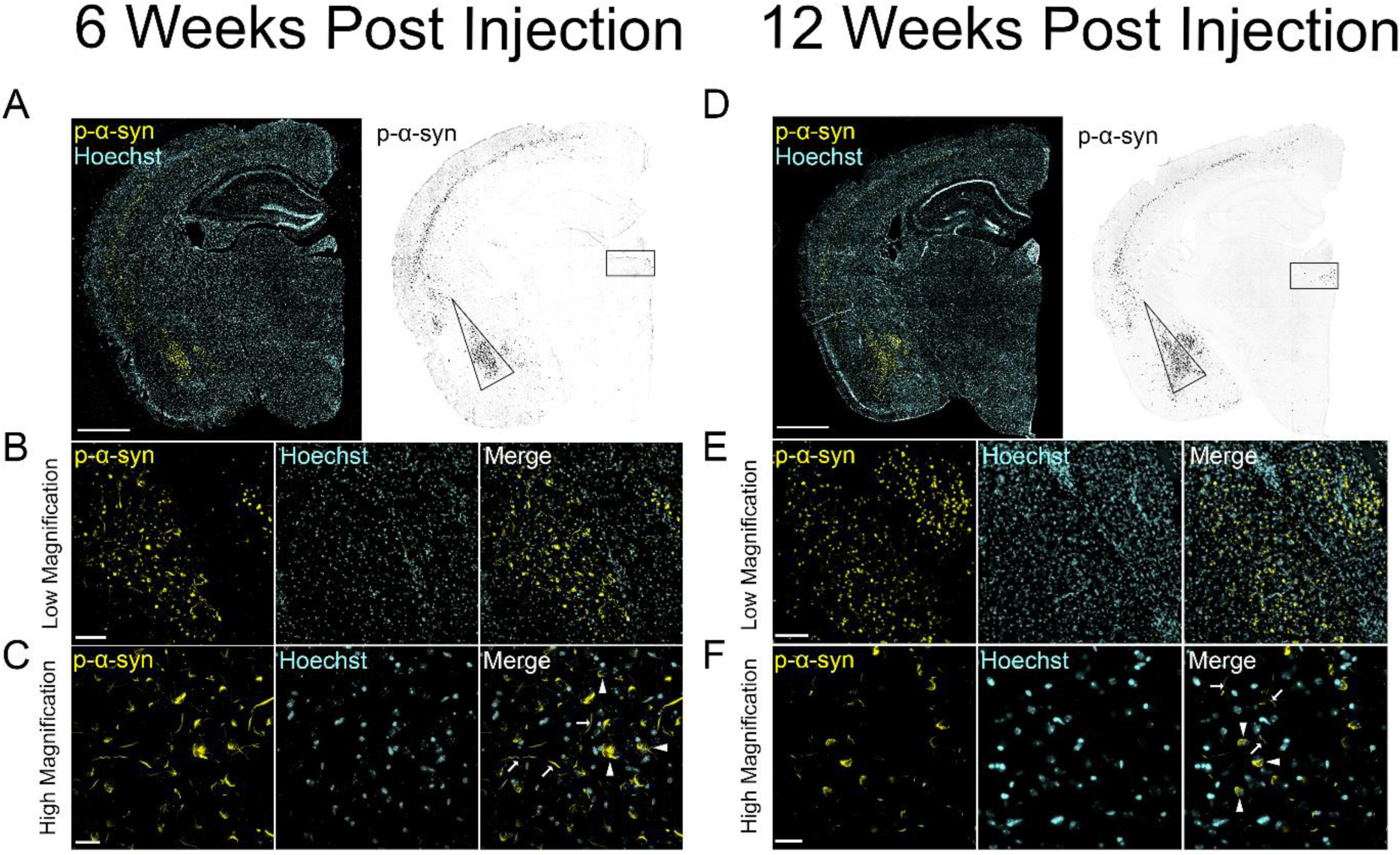
PFF injection induces α-synuclein inclusions in mouse BLA 6- and 12-weeks post-injection. Representative images for 3 – 4 month old mice injected with preformed α-syn fibrils (PFFs) and sacrificed (A - C) 6 weeks post-injection and (D - F) 12 weeks post-injection. Left panels show p-α-syn positive aggregates (yellow) and Hoechst stained nuclei (blue). (A, D) Robust p-α-syn inclusion formation observed in the basolateral amygdala (BLA), cortical layer V and the paraventricular nucleus of the thalamus shown in right panels (inverted LUT, black). (B. E) Low magnification images of p-α-syn aggregates (yellow) and nuclei (Hoechst) and (C, F) high magnification images where inclusions can be observed in the neurons of the BLA and CeA. Inclusions in the soma are indicated by white arrowheads and Lewy neurite-like inclusions indicated by white arrows. Scale Bar = 100 µm and 25 µm for low magnification and high magnification, respectively.

### Three dimensional surface reconstruction to assess changes to morphology of excitatory, cortico-amygdala synapses in mice with PFF-induced BLA α-synuclein aggregates at 6- and 12-weeks post-injection

Intrastriatal injections of PFFs cause a selective reduction in excitatory transmission of VGLUT1-positive cortico-amygdala terminals (Chen et al., 2022). To examine the effect of formation of p-α-syn aggregates on synaptic structure, we examined the morphology of synaptic puncta using excitatory, presynaptic marker VGLUT1 and excitatory, postsynaptic marker HOMER1, a scaffolding protein in excitatory post synaptic structures, at 6- and 12-weeks after intrastriatal injections of PFFs. Mice injected with phosphate buffered saline (PBS) were used as a negative control for bilateral injection into the dorsal striatum, and injection of monomeric α-synuclein was used as a negative control for the injection of protein as monomeric α-synuclein (MON) is known to not seed inclusion formation (Earls et al., 2019; Luk et al., 2012). As confirmed using immunofluorescence with an antibody to pSer129-α-syn, PBS and MON mice showed no inclusion formation when imaged with confocal at high magnification (Figure 2A) although diffuse cytosolic immunofluorescence was apparent reflecting normal p-α-syn (Froula et al., 2019; Parra-Rivas et al., 2023; Ramalingam et al., 2023). Mice injected with PFFs bilaterally into the striatum exhibited neuritic and somal inclusion formation in mouse BLA (Stoyka et al., 2020) (Figure 2A).

**Figure 2.**
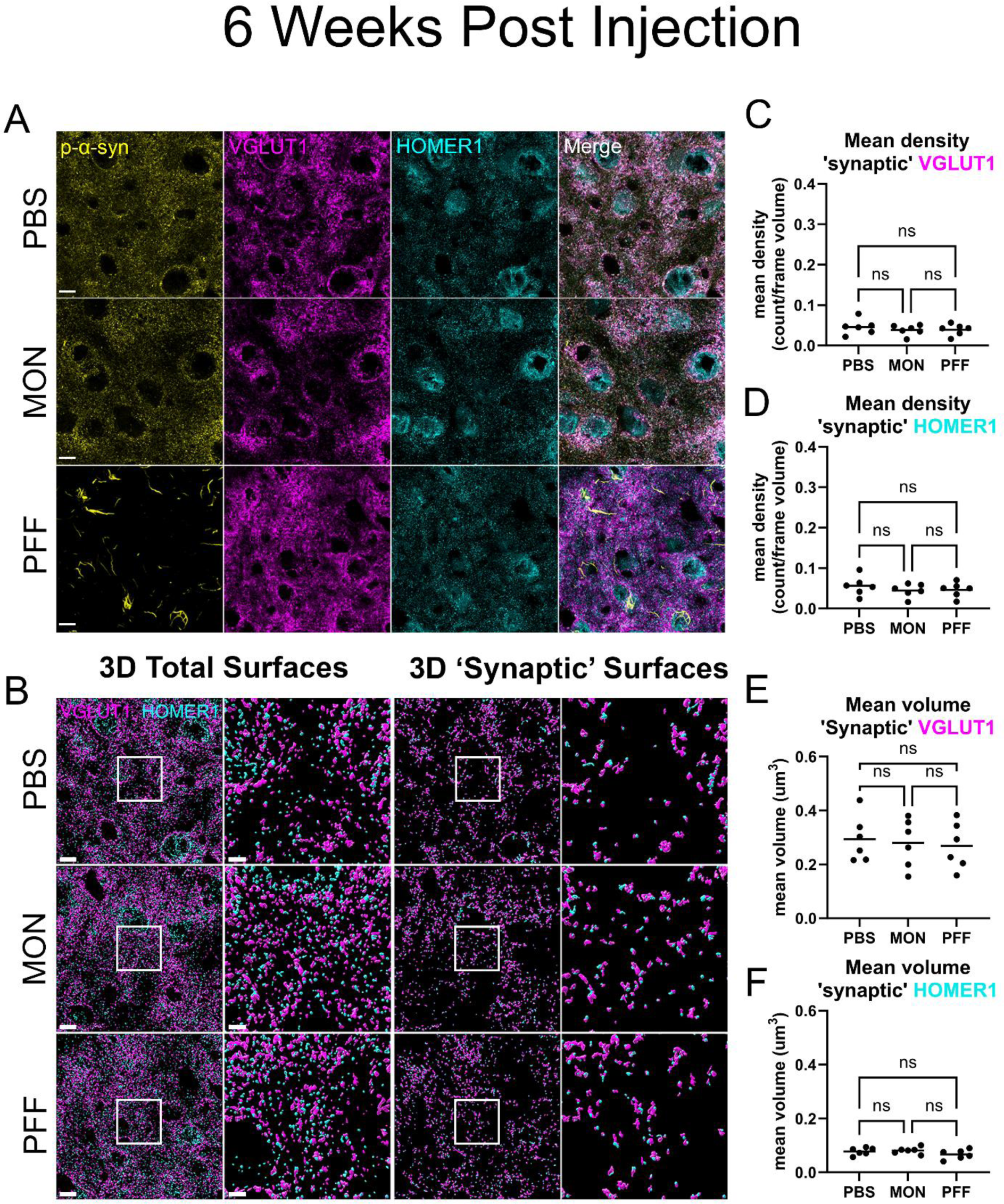
Effect of p-α-syn inclusion formation in density and volume of cortico-amygdala synaptic surfaces at 6 weeks post-injection. Mice were injected with either PBS, monomeric α–synuclein (MON), or PFFs and sacrificed 6 weeks post-injection. (A) Representative images of the deconvolved immunofluorescence for p-α-synuclein (yellow), presynaptic marker VGLUT1 (magenta) and postsynaptic marker HOMER1 (cyan). Scale bar = 10 µm. (B) 3D rendered surfaces for presynaptic VGLUT1 (magenta) and postsynaptic HOMER1 (cyan) total surfaces and 3.4X zoom inset and closely juxtaposed pre- and post-‘synaptic’ surfaces and 3.4X zoom inset. Scale bar = 10 µm and 3 µm, respectively. Mean values for the density of ‘synaptic’ surfaces for (C) VGLUT1+ and (D) HOMER1+ puncta normalized to the volume of the frame showed no significant differences between treatment and control mice. Mean values of the volume for (E) ‘synaptic’ VGLUT1+ and (F) ‘synaptic’ HOMER1+ puncta showed overall no significant differences in puncta volume for pre- and postsynaptic puncta compared to negative controls. Statistical model: One-way ANOVA with Brown-Forsythe correction applied to groups with unequal variance. Data points represent average values for each individual mouse.

Immunofluorescence was performed on glutamatergic presynaptic terminals using antibodies against presynaptic VGLUT1, and postsynaptic HOMER1 (Figure 2A). The volumes of VGLUT1 and HOMER1 puncta were rendered using Imaris 3D surface rendering software as described in methods (Figure 2B). To assess whether abnormal α-syn formation had an effect on the overall density of synaptic puncta at early time points, the number of synaptic puncta for pre- and post-synaptic surfaces was quantified 6 weeks after injection with PBS, monomeric α-syn, or PFFs. To see if there were changes to cortico-BLA synapses, the mean density per frame volume of total VGLUT1 and total HOMER1 were calculated (Supplemental Figure 3). One-way ANOVA results revealed that there were no statistically significant differences in total VGLUT1+ or total HOMER1+ mean density amongst any of the groups (Table S1A).

To assess whether abnormal α-syn formation had an effect of the number of pre- and post-synaptic puncta that are closely juxtaposed, we filtered for VGLUT1+ puncta centers that were within 0.01 μm distance from at least one HOMER1+ puncta center and vice versa. Thereafter, the number of pre- and postsynaptic puncta which were considered closely juxtaposed were counted to quantify the density of HOMER1 localized VGLUT1+ puncta/frame volume (mean density ‘synaptic’ VGLUT1) and VGLUT1 localized HOMER1+ puncta/frame (mean density ‘synaptic’ HOMER1) (Figure 2C, D). One-way ANOVA revealed there were no significant differences in ‘synaptic’ VGLUT1+ counts per frame or ‘synaptic’ HOMER1+ counts per frame (Figure 2C and D, Table 3A) between PBS, MON and PFF groups. This was taken to indicate that PFF-induced α-syn aggregate formation does not affect the density of synaptic puncta in the BLA at six weeks post-PFF injection. There was no statistical difference between the MON or PBS-injected animals.

**Table 3A.**
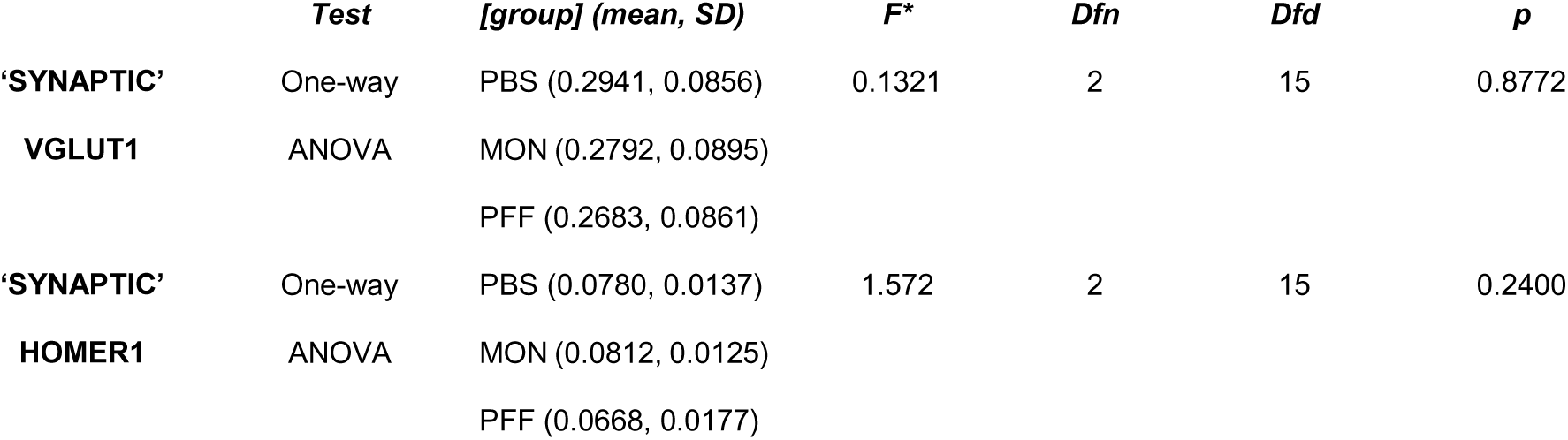
CORTICO-AMYGDALA PROJECTIONS. Statistics summary table for mean density of ‘synaptic’ VGLUT1+ and ‘synaptic’ HOMER1+ puncta 6 weeks post-injection

**Table 3B.**
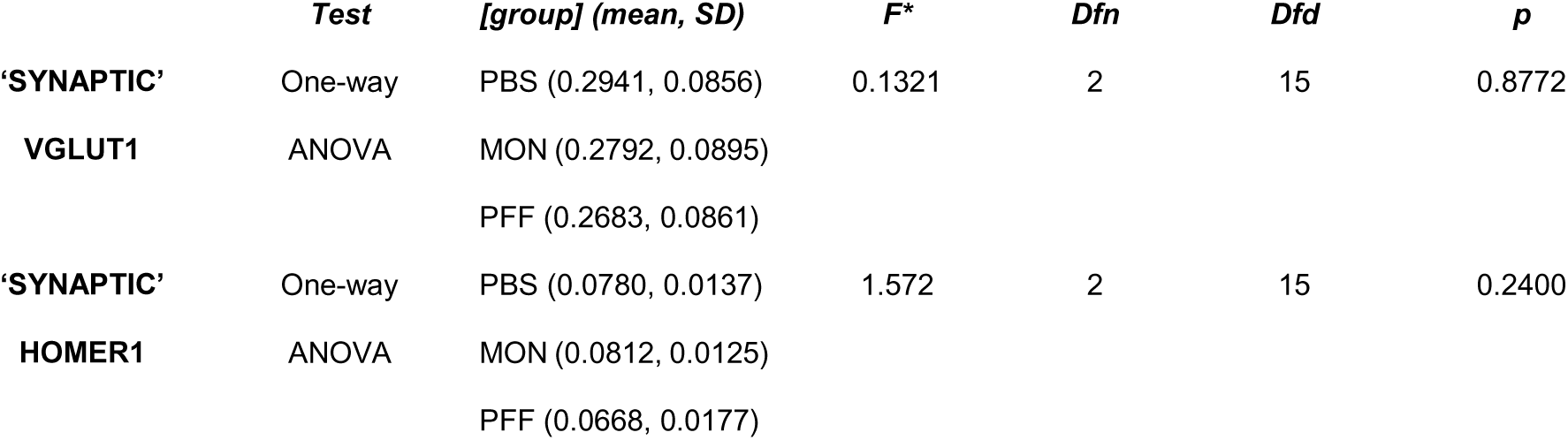
Statistics summary table for mean volume of ‘synaptic’ VGLUT1+ and ‘synaptic’ HOMER1+ puncta 6 weeks post-injection

Pathologic α-syn may cause changes in mean volume of synaptic puncta. One-way ANOVA revealed no significant differences in mean volume of total VGLUT1+ puncta between PBS-, MON-, or PFF-injected mice. However, postsynaptic puncta populations were affected by aggregate formation as mean volume of total HOMER1 positive puncta were significantly larger in PFF-injected mice. (Supplemental Figure 3, Table S1A). There was no significant difference between groups in mean volume ‘synaptic’ VGLUT1 puncta or ‘synaptic’ HOMER1 puncta (Figure 2E and F, Table 3B).

To assess if changes to cortico-amygdala synapses may occur in a time-dependent manner, we analyzed the density of VGLUT1+ and HOMER1+ puncta per frame volume for mice 12 weeks post-PFF injection (Figure 3). At 6 weeks post-injection, it was noted that there were no significant differences reported for negative controls MON and PBS-injected mice. Given that both PBS and MON do not induce α-syn inclusion formation and that there was no statistical difference between PBS and MON with respect to synapses, it was concluded that the PBS mouse cohort would be a sufficient negative control for PFF-injected mice. Hence, the remaining analyses were reported on PBS- and PFF-injected animals only.

**Figure 3.**
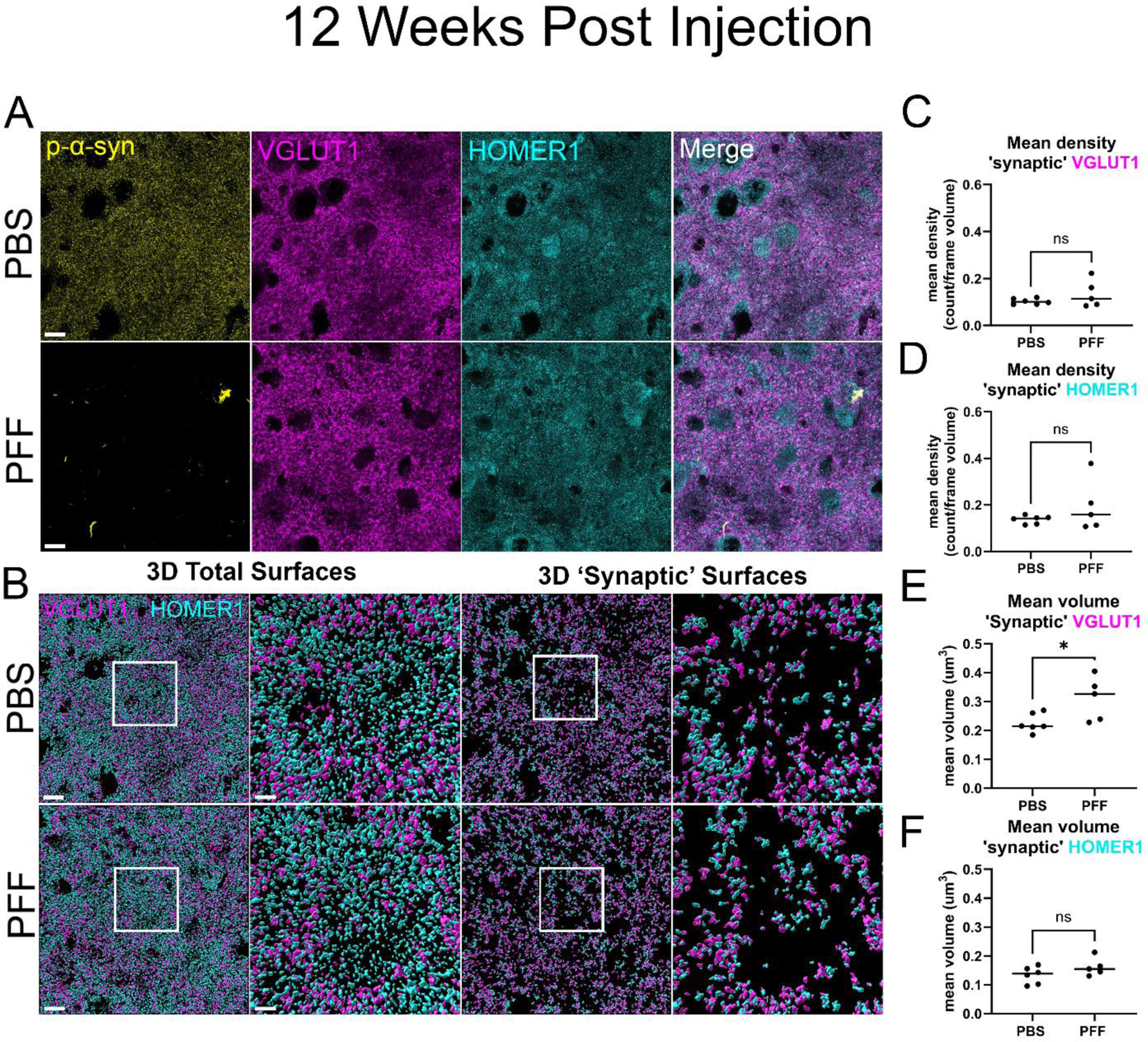
Inducing p-α-syn inclusion formation significantly increased mean volume of excitatory cortico-amygdala vGLUT1-positive terminals in mouse BLA 12 weeks post-injection. For animals injected with either PBS or PFFs 12 weeks post-injection, (A) representative images of the deconvolved immunofluorescence for p-α-syn (yellow), presynaptic marker VGLUT1 (magenta) and postsynaptic HOMER1 (cyan) signal. Scale bar = 10 µm. (B) 3D rendered surfaces for presynaptic VGLUT1 (magenta) and postsynaptic HOMER1 (cyan) total surfaces and 3.4X zoom inset and closely juxtaposed pre- and post-‘synaptic’ surfaces and 3.4X zoom inset. Scale bar = 10 µm and 3 µm, respectively. Mean values for the density of surfaces for (C) ‘synaptic’ VGLUT1+ and (D) ‘synaptic’ HOMER1+ puncta normalized to the volume of the frame showed no significant differences between treatment and control mice. Mean values of the volume for (E) ‘synaptic’ VGLUT1+ puncta showed significant increase in mean volume for PFF-injected animals compared to PBS-injected animals. (F) No significant difference in mean volume of ‘synaptic’ HOMER1+ puncta closely juxtaposed to VGLUT1+ was observed. Statistical model: Students t-test with Welch’s correction applied to groups with significant differences in variance. Data points represent average values for individual mouse.

As observed at 6 weeks post-injection, there were no significant differences in mean density for total VGLUT1+ puncta or total HOMER1+ puncta between PFF- and PBS-injected mice at 12 weeks post-injection (Supplemental Figure 4, Table S2A). For ‘synaptic’ VGLUT1+ puncta, there were no significant differences in the mean density between PBS and PFF. There were also no significant difference in the mean density of ‘synaptic’ HOMER1+ puncta between groups (Figure 3C and D, Table 4A).

**Table 4A.**
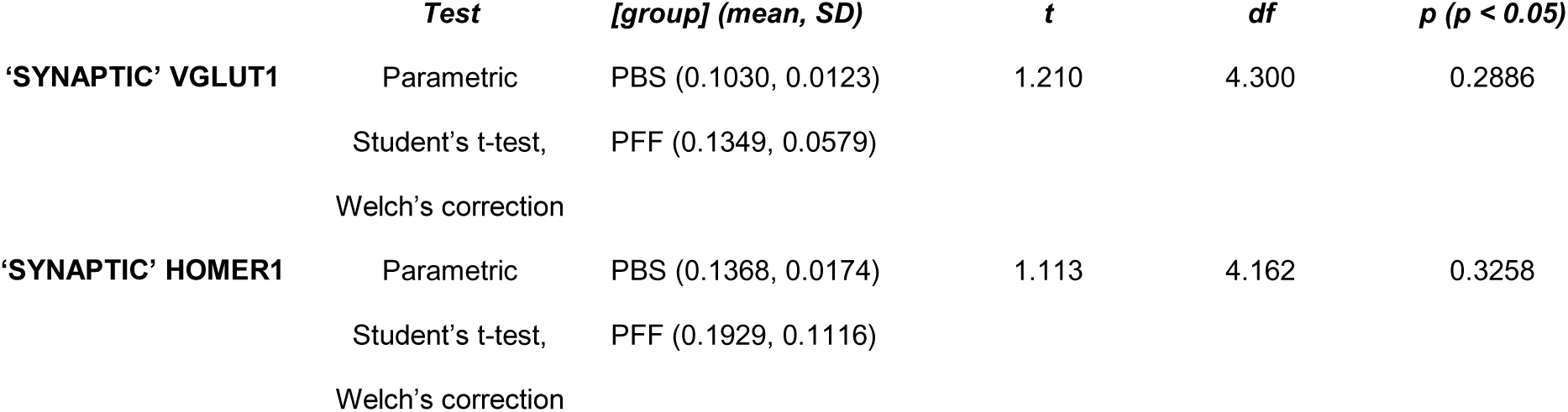
CORTICO-AMYGDALA PROJECTIONS. Statistics summary table for mean density of ‘synaptic’ VGLUT1+ and ‘synaptic’ HOMER1+ puncta 12 weeks post-injection

**Table 4B.**
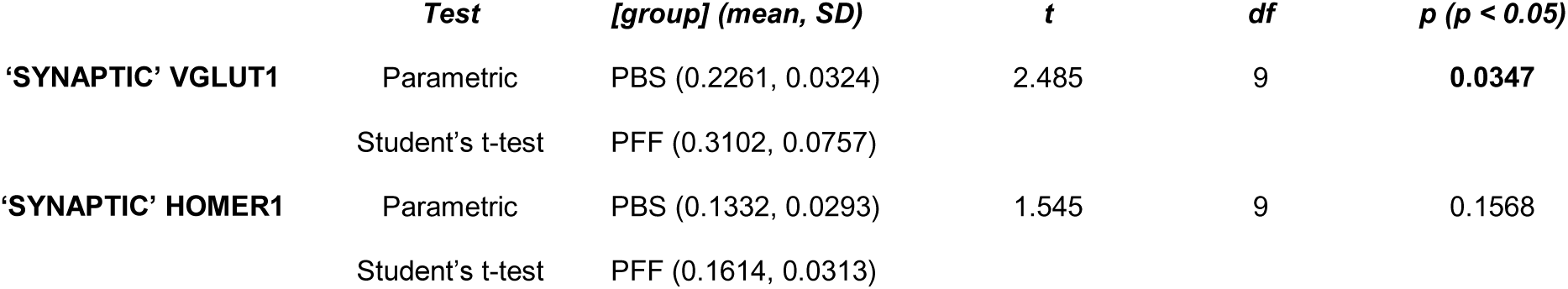
Statistics summary table for mean volume of ‘synaptic’ VGLUT1+ and ‘synaptic’ HOMER1+ puncta 12 weeks post-injection

The most robust changes in synaptic puncta were morphological as we observed significant increases in mean volume for different synaptic populations. When mean volume was measured for PFF and PBS control, we observed a significant increase in mean volume of presynaptic total VGLUT1+ puncta (Supplemental Figure 4, Table S2B). Postsynaptic puncta populations were also affected as mean volume of total HOMER1+ puncta was significantly larger in PFF-injected mice (Supplemental Figure 4, Table S2B). The mean volume of ‘synaptic’ VGLUT1+ puncta was significantly larger in PFF-injected mice than in PBS-injected mice. However, the ‘synaptic’ HOMER1+ puncta population showed no significant difference between treatment groups (Figure 3E and F, Table 4B).

### Three dimensional surface reconstruction to assess changes to morphology of excitatory, thalamo-amygdala synapses in mice with PFF-induced BLA α-synuclein aggregates at 6- and 12-weeks post-injection

To assess if VGLUT2+ thalamo-BLA synapses showed changes in density or volume, we repeated these analyses for VGLUT2+ positive terminals and HOMER1+ postsynaptic densities. To determine if there were changes to counts of thalamo-BLA terminals at 6 weeks post-injection, the mean density per frame volume of total VGLUT2+ and total HOMER1+ was calculated. There was a significant increase in mean density of total VGLUT2+ puncta per frame volume between PBS-injected mice and PFF-injected mice. There was no significant difference in mean density of total HOMER1+ puncta per frame volume between these two groups (Supplemental Figure 5, Table S3A). When synaptic puncta populations were filtered based on localization to HOMER1+ puncta, there was no significant differences in ‘synaptic’ VGLUT2+ counts per frame volume. There was also no significant difference in mean density per frame volume for ‘synaptic’ HOMER1+ (Figure 4C and D, Table 5A).

**Figure 4.**
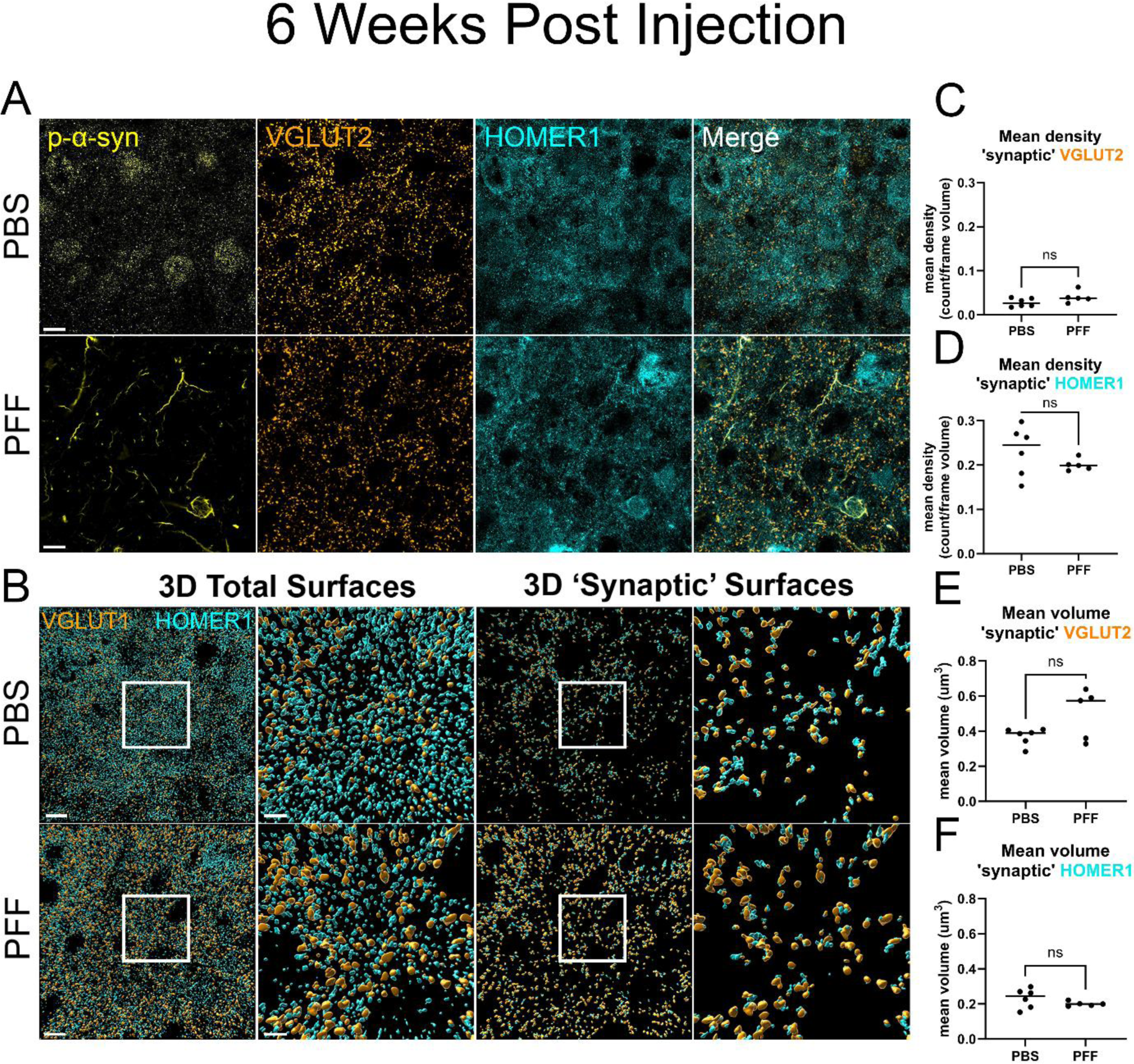
Inducing p-α-syn inclusion formation has no significant effect on density and volume of thalamo-amygdala synaptic pairs at 6 weeks post-injection. For animals injected with either PBS or PFFs 6 weeks post-injection, (A) representative images of the deconvolved immunofluorescence for p-α-syn (yellow), presynaptic marker VGLUT2 (orange) and postsynaptic HOMER1 (cyan). Scale bar = 10 µm. (B) 3D rendered surfaces for presynaptic VGLUT2 (orange) and postsynaptic HOMER1 (cyan) total surface and 3.4X zoom inset as well as closely juxtaposed pre- and post-‘synaptic’ surfaces and 3.4X zoom inset. Scale bar = 10 µm and 3 µm, respectively. Mean values for the density of ‘synaptic’ surfaces for (C) VGLUT2+ and (D) ‘synaptic’ HOMER1+ puncta normalized to the volume of the frame showed no significant differences between treatment and control mice. Mean values of the volume for (E) ‘synaptic’ VGLUT2+ and (F) ‘synaptic’ HOMER1+ puncta showed no significant differences in mean volume of puncta. Statistical model: Students t-test with Welch’s correction applied to groups with significant differences in variance. Data points represent average values for individual mouse.

**Table 5A.**
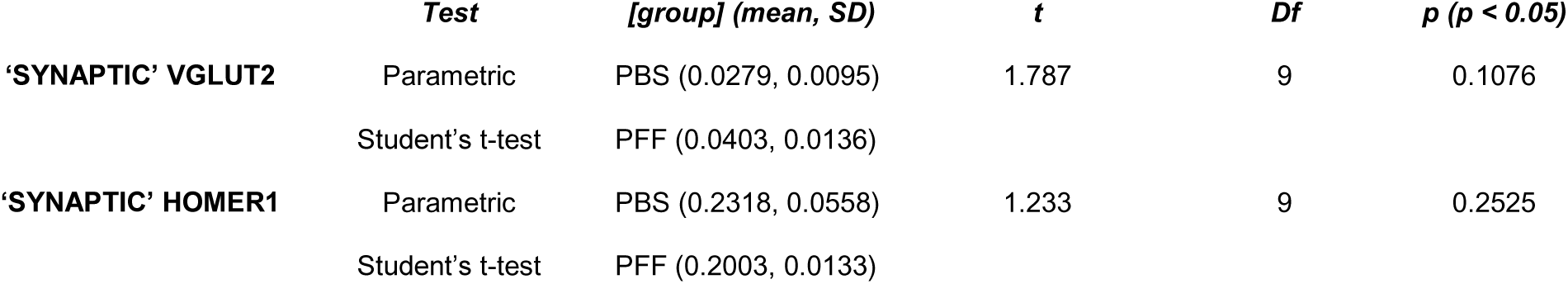
THALAMO-AMYGDALA PROJECTIONS. Statistics summary table for mean density of ‘synaptic’ VGLUT2+ and ‘synaptic’ HOMER1+ puncta 6 weeks post-injection

**Table 5B.**
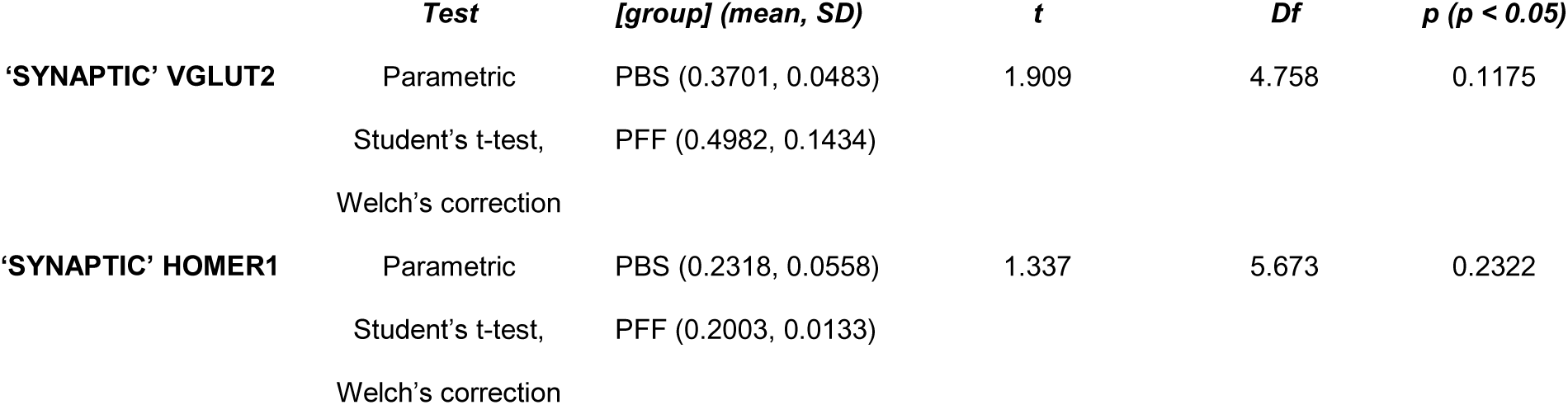
Statistics summary table for mean volume of ‘synaptic’ VGLUT2+ and ‘synaptic’ HOMER1+ puncta 6 weeks post-injection

To assess if changes to thalamo-BLA projections may affect the morphology of synaptic puncta, we analyzed the mean volume of total VGLUT2+ and total HOMER1+ puncta. There was no significant difference in mean volume of total VGLUT2+ puncta between PBS- and PFF-injected mice with mean volumes of PFF-injected mice. We observed no significant differences in mean volume of total HOMER1+ puncta between PBS-injected mice and PFF-injected mice (Supplemental Figure 6, Table S3B).

When surfaces were filtered to isolate closely juxtaposed populations, no changes were observed in the mean volume of ‘synaptic’ puncta. When isolating ‘synaptic’ VGLUT2+ puncta, we observed no significant differences in mean puncta volume between PBS-injected mice and PFF-injected mice. Similarly, we report no significant difference in mean volume of VGLUT2 localized, ‘synaptic’ HOMER1+ puncta between PBS-injected mice and PFF-injected mice (Figure 4E and F, Table 5b).

To see if changes to thalamo-BLA projection counts might be time-dependent, the mean density per frame volume of total VGLUT2+ and total HOMER1+ at 12-weeks-post-injection were also calculated. There were no significant differences in mean density of total VGLUT2+ puncta per frame volume between PBS-injected mice and PFF-injected mice. There were also no significant differences in mean density per frame volume of total HOMER1+ puncta between PBS-injected mice and PFF-injected mice (Supplemental Figure 6, Table S4A). When presynaptic VGLUT2+ puncta populations were filtered based on localization to HOMER1+ puncta, there were no significant differences in ‘synaptic’ VGLUT2+ counts per frame volume between groups. There were also no significant differences in mean density per frame volume for ‘synaptic’ HOMER1+ puncta between PBS- and PFF-injected mice (Figure 5C and D, Table 6A). Overall, we observed no significant differences in mean density of thalamo-amygdala VGLUT2+/HOMER1+ projections between PBS- and PFF-injected animals at 12 weeks post-injection.

**Figure 5.**
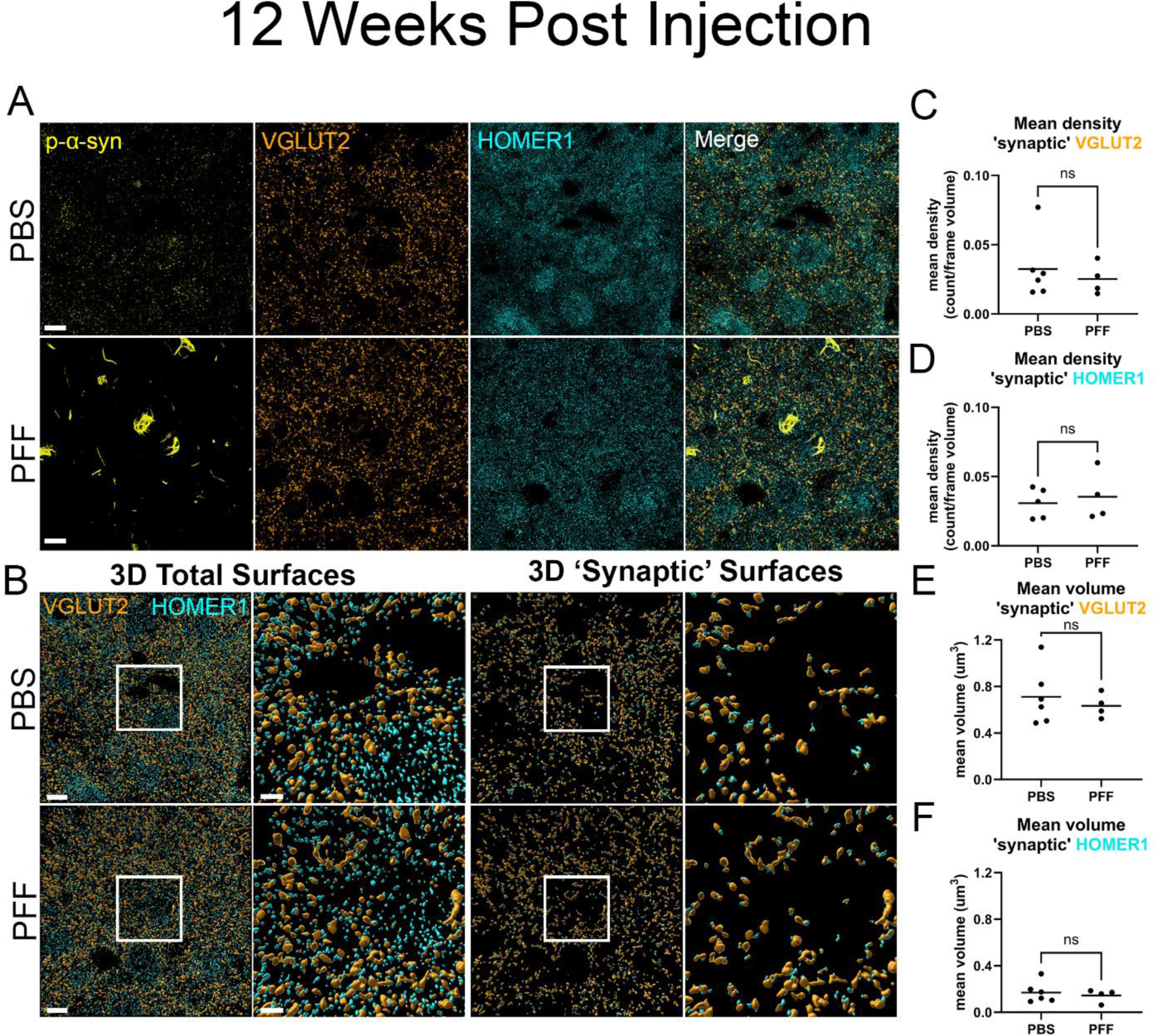
Inducing p-α-syn inclusion formation has no significant effect on density and volume of thalamo-amygdala projections 12 weeks post-injection. (A) For animals inoculated with either PBS or PFFs 12 weeks post-injection, representative images of the deconvolved immunofluorescence for p-α-syn (yellow), presynaptic marker VGLUT2 (orange) and postsynaptic HOMER1 (cyan). (B) 3D rendered surfaces for presynaptic VGLUT2 (orange) and postsynaptic HOMER1 (cyan) total surfaces and 3.4X zoom inset and closely juxtaposed pre- and post-‘synaptic’ surfaces and 3.4X zoom inset. Scale bar = 10 µm and 3 µm, respectively. Mean values for the density of (C) ‘synaptic’ VGLUT2+ and (D) ‘synaptic’ HOMER1+ puncta normalized to the volume of the frame showed no significant differences between treatment and control mice. Mean values of the volume for (E) ‘synaptic’ VGLUT2+ and (F) ‘synaptic’ HOMER1+ puncta showed no significant differences in mean volume of puncta. Statistical model: Students t-test with Welch’s correction applied to groups with significant differences in variance (p > 0.5). Data points represent average values for individual mouse.

**Table 6A.** THALAMO-AMYGDALA PROJECTIONS. Statistics summary table for mean density of ‘synaptic’ VGLUT2+ and ‘synaptic’ HOMER1+ puncta 12 weeks post-injection

|  | *Test* | *[group] (mean, SD)* | *T* | *df* | *p (p < 0.05)* |
| --- | --- | --- | --- | --- | --- |
| 'SYNAPTIC' VGLUT2 | Parametric | PBS (0.0325, 0.0228) | 0.5747 | 8 | 0.5813 |
|  | Student's t-test | PFF (0.0253, 0.0113) |  |  |  |
| 'SYNAPTIC' HOMER1 | Parametric | PBS (0.0308, 0.0108) | 0.4836 | 7 | 0.6434 |
|  | Student's t-test | PFF (0.0354, 0.0178) |  |  |  |

**Table 6B.** Statistics summary table for mean volume of ‘synaptic’ VGLUT2+ and ‘synaptic’ HOMER1+ puncta 12 weeks post-injection

|  | *Test* | *[group] (mean, SD)* | *T* | *df* | *p (p < 0.05)* |
| --- | --- | --- | --- | --- | --- |
| 'SYNAPTIC' VGLUT2 | Parametric | PBS (0.7108, 0.2433) | 0.5884 | 8 | 0.5725 |
|  | Student's t-test | PFF (0.6339, 0.1032) |  |  |  |
| 'SYNAPTIC' HOMER1 | Parametric | PBS (0.1701, 0.0886) | 0.5161 | 8 | 0.6197 |
|  | Student's t-test | PFF (0.1442, 0.0554) |  |  |  |

To assess if morphological changes in thalamo-BLA synapses may be time-dependent, we also assessed the mean volume of total VGLUT2+ and total HOMER1+ puncta at 12-weeks-post-injection. There was no significant difference in mean volume of total VGLUT2+ puncta between PBS- and PFF-injected mice. We also observed no significant differences in mean volume of total HOMER1+ between PBS-injected mice and PFF-injected mice (Supplemental Figure 6, Table S4B). There were also no changes observed in the mean volume of ‘synaptic’ puncta when surfaces were filtered to isolate closely juxtaposed populations. ‘Synaptic’ VGLUT2+ puncta showed no significant differences in mean puncta volume between PBS-injected mice and PFF-injected mice. Similarly, we report no significant difference in mean volume of ‘synaptic’ HOMER1+ puncta between PBS-injected mice and PFF-injected mice (Figure 5E and F, Table 6B). Thus, while cortico-BLA, VGLUT1/HOMER1 synapses in the BLA show morphological changes at 12 weeks post-PFF injections, VGLUT2/HOMER1 thalamo-BLA synapses do not.

### Three-dimensional surface reconstruction to assess changes to morphology of BLA excitatory synapses containing α-syn aggregates at 6- and 12-weeks post-injection

We observed that although p-α-syn inclusion formation in the BLA is robust, only a small proportion of synapses are also positive for p-α-syn aggregates, and some puncta contain aggregates that appear as small puncta. It was previously shown that in DLB cortex, presynaptic terminals containing p-α-syn show increased volumes (Colom-Cadena et al., 2017). We hypothesized that the presence of p-α-syn within synaptic puncta would have an effect on the overall morphology of those puncta and that those changes may be absent in those puncta that do not have p-α-syn inclusions.

Total VGLUT1+ puncta were separated into VGLUT1 with p-α-syn and without p-α-syn. The mean volume of total VGLUT1+ puncta colocalized with p-α-syn (w p-α-syn) was significantly higher than that of VGLUT1+ puncta that were not colocalized with p-α-syn (wo p-α-syn) at 6-weeks post-injection. Similarly, postsynaptic puncta were separated to assess differences in total HOMER1+ puncta with and without p-α-syn inclusions. Again total HOMER1+ mean volume was significantly higher for total HOMER1+ puncta containing p-α-syn compared to total HOMER1 not containing p-α-syn (Figure 6A and C, Table 7A).

**Figure 6.**
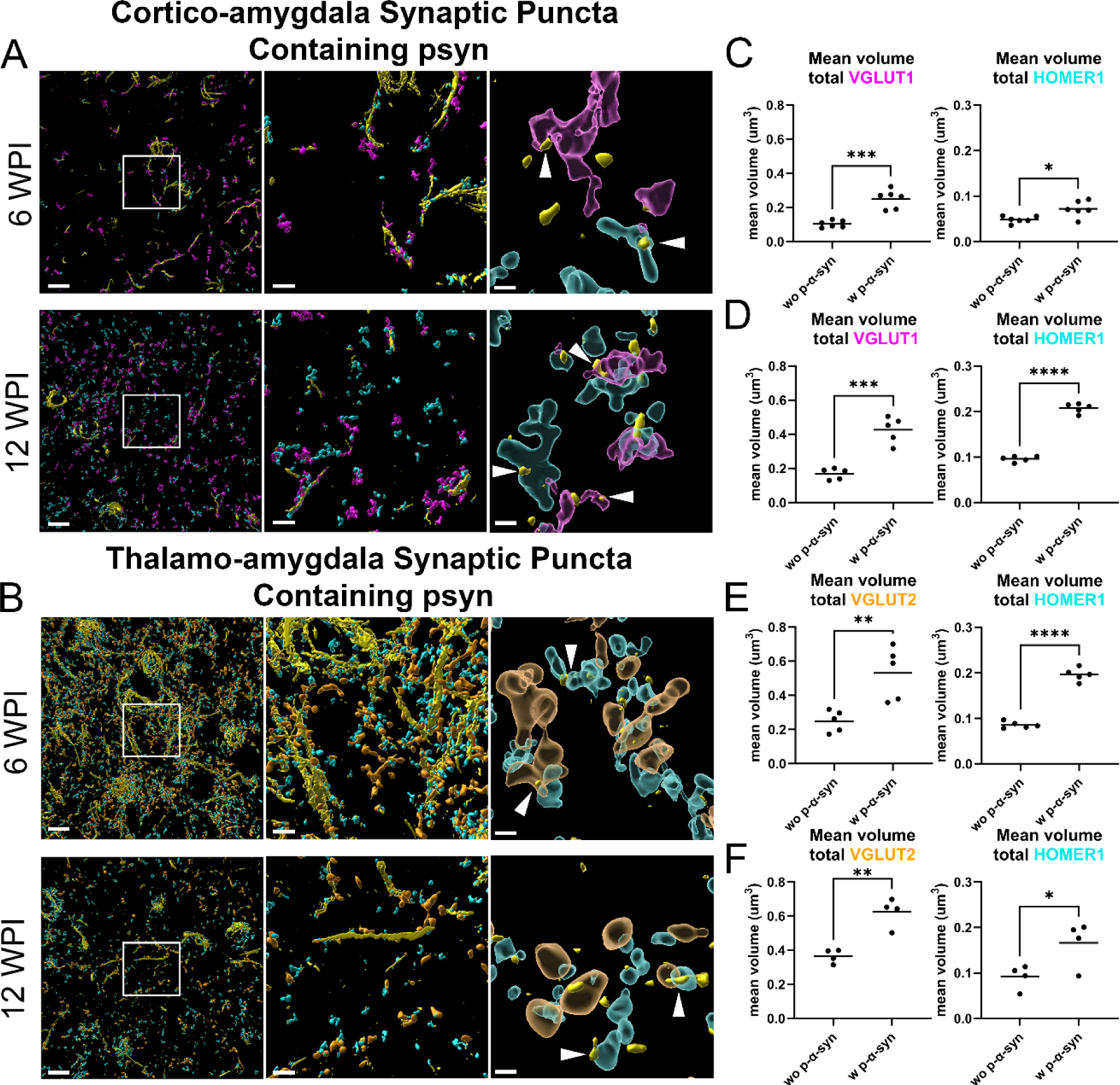
Volume of synaptic puncta containing p-α-syn are larger than those that do not contain p-α-syn aggregates at 6- and 12-weeks post-injection. For animals injected with PFFs, representative images of 3D rendered surfaces for p-α-syn inclusions and pre- and postsynaptic excitatory markers closely juxtaposed to inclusions. (A) 3D surfaces for VGLUT1+ (magenta) and HOMER1+ (cyan) cortico-amygdala projections containing p-α-synuclein inclusions (yellow). (B) 3D surfaces for VGLUT2+ (orange) and HOMER1+ (cyan) thalamo-amygdala projections containing p-α-synuclein inclusions (yellow). Second inset represents only synaptic puncta containing p-α-syn aggregates. For visual clarity, p-α-syn aggregates with volume greater than 0.1 µm have been removed. White arrows used to indicate synaptic puncta containing p-α-syn aggregates with volume below 0.1 µm. Scale bar left = 10 µm, scale bar middle = 3 µm & scale bar right = 0.5 µm. (C – F) Mean volume of pre- and postsynaptic puncta show significant increase in mean volume of puncta when those puncta are positive for p-α-synuclein (w p-α-syn) compared to those puncta without p-α-synuclein (wo p-α-syn) measured as distance from surface between synaptic puncta and p-α-synuclein for cortico-amygdala (VGLUT1/HOMER1) and thalamo-amygdala (VGLUT2/HOMER1) projections. Statistical model: Students t-test with Welch’s correction applied to groups with significant differences in variance. Data points represent average values for individual mouse.

**Table 7A.** CORTICO-AMYGDALA PROJECTIONS. Statistics summary table for mean total volume of total VGLUT1+ and total HOMER1+ puncta containing p-α-syn 6 weeks post-injection

|  | *Test* | *[group] (mean, SD)* | *t* | *df* | *P (p < 0.05)* |
| --- | --- | --- | --- | --- | --- |
| **TOTAL VGLUT1** | Parametric | wo p-α-syn (0.1036, 0.0206) | 6.188 | 10 | **0.0001** |
| | Student's t-test | w p-α-syn (0.2507, 0.0545) | | | |
| **TOTAL HOMER1** | Parametric | wo p-α-syn (0.0484, 0.0076) | 2.981 | 10 | **0.0138** |
| | Student's t-test | w p-α-syn (0.0720, 0.0178) | | | |

**Table 7B.** Statistics summary table for mean volume of total VGLUT1+ and total HOMER1+ puncta containing p-α-syn 12 weeks post-injection

|  | *Test* | *[group] (mean, SD)* | *t* | *df* | *P (p < 0.05)* |
| --- | --- | --- | --- | --- | --- |
| **TOTAL VGLUT1** | Parametric | wo p-α-syn (0.1693, 0.0325) | 6.910 | 8 | **0.0001** |
| | Student's t-test | w p-α-syn (0.4267, 0.0767) | | | |
| **TOTAL HOMER1** | Parametric | wo p-α-syn (0.0962, 0.0063) | 20.47 | 8 | **<0.0001** |
| | Student's t-test | w p-α-syn (0.2077, 0.0104) | | | |

**Table 7C.**
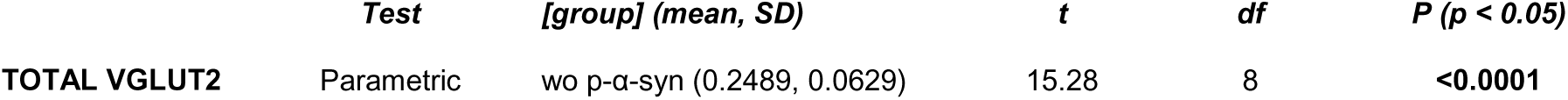
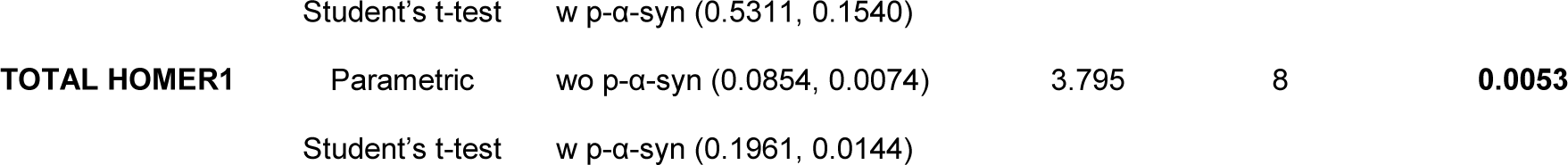
THALAMO-AMYGDALA PROJECTIONS. Statistics summary table for mean volume of total VGLUT2+ and total HOMER1+ puncta containing p-α-syn 6 weeks post-injection

**Table 7D.**
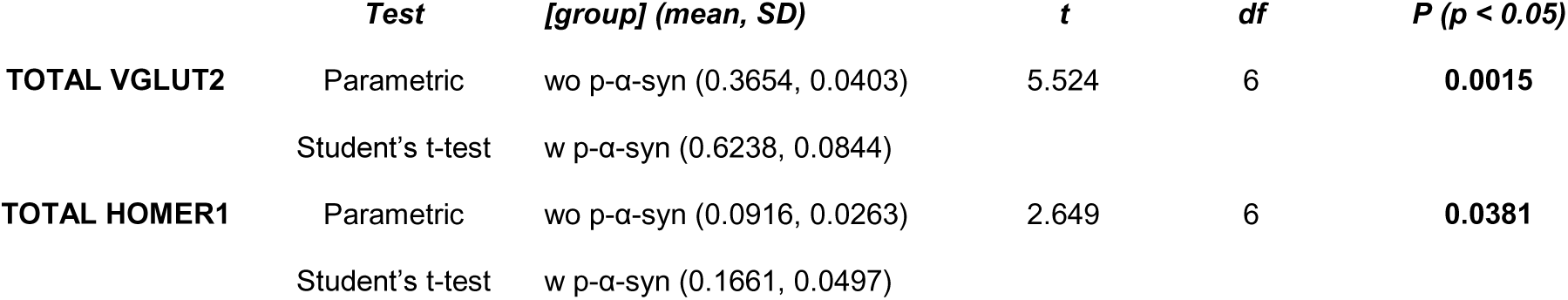
Statistics summary table for mean volume of total VGLUT2+ and total HOMER1+ puncta containing p-α-syn 12 weeks post-injection

To determine if the effect of inclusions persisted with time, we repeated this analysis with PFF-injected mice at 12-weeks post-injection. As seen at earlier time points, the mean volume of total VGLUT1+ puncta colocalized with p-α-syn was significantly higher than that of total VGLUT1+ puncta not colocalized with p-α-syn. This effect was also found in postsynaptic puncta as mean volume of total HOMER1+ puncta containing p-α-syn was significantly higher than that of total HOMER1+ puncta not containing p-α-syn (Figure 6A and D, Table 7B).

To see if there was a similar effect on mean volume of thalamo-amygdala synapses containing p-α-syn, this analysis was performed on 6- and 12-week post-injection cohorts of PFF mice. After separating surface populations into total VGLUT2+ puncta with p-α-syn and without p-α-syn and total HOMER1+ puncta with p-α-syn and without p-α-syn as already described, we compared the mean volume of these populations. At 6-weeks post-injection, the mean volume of total VGLUT2+ puncta with p-α-syn was significantly higher than without p-α-syn. Similarly, the mean volume of total HOMER1+ puncta with p-α-syn was also significantly higher than that of total HOMER1+ puncta without p-α-syn (Figure 6B and E, Table 7C).

Similar to the findings for cortico-amygdala projections, it was found that the effect on synaptic volumes when those puncta contained p-α-syn was consistent at 12-weeks-post-injection. For this cohort of mice, the mean volume of total VGLUT2+ puncta colocalized with p-α-syn was significantly higher than mean volume of total VGLUT2+ puncta not colocalized with p-α-syn. Postsynaptic populations were affected similarly with mean volume of total HOMER1+ puncta containing p-α-syn being significantly higher than the mean volume of total HOMER1+ puncta not containing p-α-syn (Figure 6B and F, Table 7d). Overall, the morphology of cortico-amygdala projections as well as thalamo-amygdala projections is significantly affected by the presence of p-α-synuclein aggregates within synaptic puncta.

### Transmission electron microscopy to assess ultrastructural changes to excitatory synapses in mice with PFF-induced BLA α-syn aggregates

Transmission electron microscopy was performed to determine if there are changes in synaptic morphology in the BLA of mice that received intrastriatal injections of PBS or PFFs. Mice were analyzed 12 weeks post-injections. After perfusions, the BLA was dissected, and TEM was performed. For analyses, asymmetric, excitatory synapses were identified and the PSD length and number of docked vesicles per PSD length were measured. The area of individual synaptic vesicles and the nearest neighbor distance of synaptic vesicles were measured using a published convolutional neural network algorithm trained on mouse synapses (Imbrosci et al., 2022) (Figure 7). In the BLA, there were no significant differences between PBS- and PFF-injected mice for length of PSD (Figure 7B), number of docked vesicles normalized to PSD length (Figure 7C). The length of the synaptic boutons were also measured and although the data showed a bimodal distribution, there were no differences between PBS- and PFF-injected mice (Supplemental Figure 7A). There were, however, significant differences in the inter-vesicular distances. The mice with PFF-injections show reduced distance from one synaptic vesicle to its nearest neighbor (Figure 7D). The neural network algorithm used thresholds for the synaptic vesicle area measurements and the data did not appear continuous (Supplemental Figure 7B). Therefore, the synaptic vesicle areas were analyzed by comparing the percentage of cases that were above or below the grand median for PBS and PFF. There were statistically significant differences, and the data showed that in PFF-injected mice, there is a lower percentage of “larger” synaptic vesicle areas and a higher percentage of “smaller” synaptic vesicle areas. (Figure 7E). The synaptic vesicles appeared more compact within the terminal, similar to previous findings in hippocampal neurons exposed to PFFs (Froula et al., 2018).

**Figure 7.**
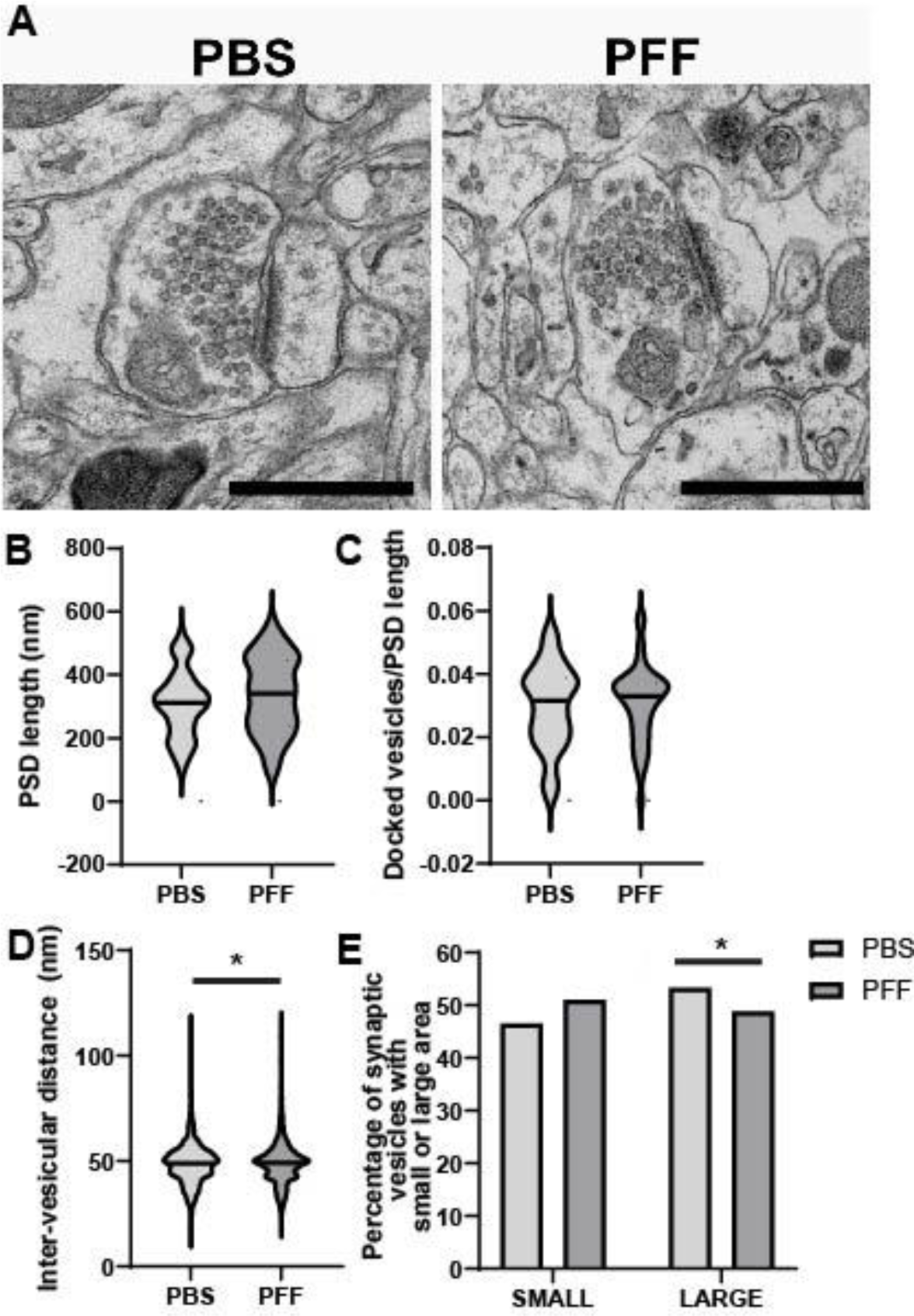
Intervesicular distances and synaptic vesicle volumes are reduced in excitatory synapses in the BLA of PFF-injected mice. At 12 weeks post-injection, transmission electron microscopy was performed using the BLA from PBS (N=4) and PFF (N=4) mice. **(A)** Representative image of asymmetric synapses quantified from PBS- and PFF-injected mice (Scale bar = 600 nm). (B) Fiji was used to quantify the length of the PSD in asymmetric synapses (n=412). Data were log10 transformed and analyzed using linear mixed models with synapses nested within each mouse, treatment as a fixed variable and compound symmetry as the covariance type. (F(1,6)=1.7, p = 0.245). (C) 37.8, p < 0.001. Docked vesicles were defined as SVs the fell within distance ≤ 100 nm from active zone adjacent to traced PSD length. The number of docked vesicles was divided by the PSD length measured for that synapse. Data were log10 transformed and analyzed using linear mixed models with synapses nested within each mouse, treatment as a fixed variable and compound symmetry as the covariance type. (F(1,6)= 0.15, p = 0.71). (D, E) Intervesicular distances and synaptic vesicle were measured using a previously published convolution neural network algorithm (n=10,400 for PBS, n=9,570 for PFF). For (D), intervesicular distances, data were analyzed using linear mixed models with synapses nested within each mouse, treatment as fixed variable and compound symmetry as the covariance type. (F(1,6)=14.3, p = 0.007). For (E), the synaptic vesicle areas did not fit a continuous distribution because of the thresholding built in to the neural network. The data were thus binned into cases above or below the grand median for PBS and PFF. A Fisher’s exact test revealed significantly higher percentage of smaller synaptic vesicle areas in the PFF treated group χ^2^ = 37.9, p<0.001).

## Discussion

Lewy pathology is found in multiple brain regions in Parkinson’s disease and Dementia with Lewy bodies. It is particularly common and abundant in the amygdala where it could contribute to cognitive and psychiatric symptoms. The intrastriatal PFF model induces formation of α-syn aggregates in the cortex and amygdala, particularly in excitatory neurons (Stoyka et al., 2020). Mice with PFF-induced α-synuclein aggregates show behavioral defects associated with amygdala function amygdala, as well as robust deficits in cortico-amygdala transmission (Chen et al., 2022). The goal of this study was to determine if changes occur in excitatory synapse morphology and density at early and later time points after initiation of α-synuclein aggregation. At 6- and 12-weeks post-PFF injections, the BLA shows abundant α-synuclein inclusions. However, the density of both cortico-amygdala and thalamo-amygdala synapses was unchanged at both time points. The most robust finding was a significant increase in the volume of pre-synaptic terminals and post-synaptic volumes in both cortico-amygdala and thalamo-amygdala synapses harboring p-α-syn. Analyses of synapse ultrastructure revealed more densely packed synaptic vesicles in PFF-injected animals compared to PBS controls at 12 weeks post-injection as well as a greater percentage of synaptic vesicles with a small area.

Increased volume of synapses in α-synucleinopathies and Parkinson’s disease models has now been demonstrated in a few studies, thus supporting our findings as a bona fide morphological change. In the DLB post-mortem cingulate cortex, presynaptic terminals with small aggregates of p-α-syn showed increased volumes (Colom-Cadena et al., 2017). Both cortico-striatal and thalamo-striatal terminals and postsynaptic densities show increases in volume in non-human primates treated with 1-methyl-4-phenyl-1,2,3,6-tetrahydropyridine as a model of Parkinson’s disease (Villalba & Smith, 2011). The mechanisms by which synapses show increased volumes are is yet unknown. Synucleins play a role in endocytosis of synaptic vesicles and thus sequestration of normal α-syn into aggregates could cause a loss of function phenotype, reducing retrieval of exocytosed synaptic vesicles, leading to expansion of the presynaptic plasma membrane (Vargas et al., 2014). Another mechanism could be expansion of organelles such as endoplasmic reticulum or mitochondria in an attempt to protect synapses from toxic α-syn aggregates. Increased synaptic activity could also cause an increased synaptic volume. Although it has been shown that PFF-induced aggregates in the amygdala cause an overall reduction in cortico-striatal transmission (Chen et al., 2022), this study was not able to specifically record from synapses harbouring small α-syn aggregates because of the current lack of tools to identify them in live, unfixed neurons. In the future, the development of tools to identify synapses with small aggregates live could better help determine the relationships between synaptic transmission, morphology and presence of p-α-syn.

It has recently been shown that phosphorylation of native α-synuclein is physiologically necessary for activity-dependent association with synapsin and VAMP2 (Parra-Rivas et al., 2023). This again brings into question whether the sequestration of α-syn into these aggregates not only reduces the level of native α-synuclein available within presynapses but also prevents α-syn from phosphorylating and dephosphorylating as necessary to facilitate binding and vesicle release at the level necessary to maintain synapse function. Transmission electron microscopy was performed to determine if mice with PFF-induced α-syn aggregates in the amygdala show changes in synaptic morphology or location of synaptic vesicles. This technique is limited because it was not possible to identify synapses with p-α-syn aggregates or distinguish cortico-amygdala or thalamo-amygdala synapses. It is possible that TEM analysis of synapses that specifically contain inclusions would show differences in synaptic vesicles in that population of synapses compared to controls or synapses without these inclusions as seen using immunofluorescence. Overall, there were no changes in length of postsynaptic density or synaptic boutons. The number of docked vesicles was also not changed, indicating no changes in the readily releasable pool of synaptic vesicles. The TEM did show reduced distance between individual synaptic vesicles as was seen in primary hippocampal neurons exposed to PFFs (Froula et al., 2018). A similar clustering of vesicles also occurs in neurons lacking all isoforms of synuclein (Vargas et al., 2014) and in neurons expressing phospho-mimetic α-syn S129D (Parra-Rivas et al., 2023). In addition, expression of phospho-mimetic α-syn S129D attenuates synaptic vesicle recycling. Chen et al found that PFF-injection mediates a reduction in frequency but not amplitude of optogenetically evoked, cortico-BLA asynchronous Sr2^+^-induced EPSCs. Additionally, they found no changes in density of VGLUT1+ terminals and no changes in initial release probability leading them to conclude that fewer synaptic vesicle release sites may explain reduction in cortico-BLA transmission (Chen et al., 2022). Thus, the formation of small p-α-syn aggregates could be a mechanism for the loss of cortico-amygdala transmission and specifically related to changes in the presynaptic compartment.

The increased volume in synapses with p-α-syn aggregates could be a homeostatic response to the formation of pathologic aggregates. A recent proteomic study of the amygdala from Parkinson’s disease brains indicates that there may be some measures in place to protect the balance of synaptic proteins in the amygdala complex of PD patients. Compared to controls, PD amygdala showed upregulation of NPTN, which has been reported in induction of neurite outgrowth and regulation of synaptic structure, function and plasticity (Beesley et al., 2014; Villar-Conde et al., 2023; J. Zhang et al., 2021; Zhang et al., 2022). This indicates that a protective phenotype may be active in PD to maintain and retain synapses.

Previous studies have shown that an abundance of α-synuclein micro-aggregate accumulations at the presynaptic terminal correlate with a downregulation in presynaptic proteins, such as syntaxin and synaptophysin, as well as postsynaptic proteins, PSD95 and drebrin (Kramer & Schulz-Schaeffer, 2007). Formation of α-syn aggregates have also been shown to correspond with major loss of dendritic spines in the mouse primary hippocampal culture models, and in the cortex *in vivo,* using PFF and in α-syn overexpression mouse models (Blumenstock et al., 2017; Froula et al., 2018). We were not able to recapitulate these findings as our study did not show changes in the density of pre- and postsynaptic puncta in the BLA at 6- and 12-weeks post-injection. It is worth noting that while mushroom spines in cultured neurons are thought to represent more mature excitatory synapses, the synaptic integrity and complex circuitry represented by *in vivo* analysis within the BLA may contribute to major differences in analysis outcomes. The effects of plasticity and other circuit effects such as inhibitory tone and changes in protein expression may mitigate the toxic effects of p-α-syn pathology as measured in cultured neurons. In the brain, a loss of spines in the cortex with no apparent loss of excitatory synapses in the BLA may reflect regional differences in the chemical and electrical environments between these two brain regions. Interestingly, in layer I of the cortex of PFF-injected mice, there is an increase in density of stubby spines which are typically found to be more dense in post-natal brains (Blumenstock et al., 2017). Overall, our data point to a possible homeostatic changes in synaptic morphology at earlier time points after initiation of p-α-syn aggregation.

The abundance of literature aimed at elucidating the changes in synaptic abundance, synaptic protein homeostasis and synaptic function in the PD brain indicates that the question of how synapses are changing in relation to α-synuclein aggregation is of special relevance to the field. Understanding the dynamics of synapse morphology and the changes to the organization of synaptic vesicles in glutamatergic synapses is vital to devise effective therapeutic strategies to protect these synapses in the brain especially given the vulnerability of glutamatergic synapses to α-synuclein aggregation in the PD brain. Studying the BLA could equip scientists with the information to understand the vulnerabilities of different glutamatergic populations that could relate to the overall glutamatergic network in the PD brain.

## Abbreviations

α-syn: α-synuclein
BLA: basolateral amygdala
DLB: Dementia with Lewy Bodies
MON: monomeric α-synuclein
MRI: Magnetic Resonance Imaging
NPTN: neuroplastin
PD: Parkinson’s Disease
p-α-syn: phosphorylated-α-synuclein
PBS: phosphate buffered saline
PET: positron emission tomography
PFA: paraformaldehyde
PFFs: preformed fibrils
PSD: post synaptic density
PSD95: post synaptic density protein 95
SNc: substantia nigra pars compacta
SVs: synaptic vesicles
TBS: tris-buffered saline
TEM: transmission electron microscopy
VGLUT: vesicular glutamate transporter
w p-α-syn: with phophorylated-α-synuclein
wo p-α-syn: without phosphorylated-α-synuclein

## Acknowledgements

We would also like to thank the UAB High Resolution Imaging Facility led by Dr. Alexa Mattheyses who was on Nolwazi Gcwensa’s thesis committee and provided expertise and guidance on this project. We are grateful to Dr. Jacqueline Burre of Weill Cornell Medical School for sharing her protocol for perfusing mice for electron microscopy. This work was supported by the National Institutes of Health, NINDS grant R56NS117465, American Parkinson Disease Association grant 977962, Aligning Science Across Parkinson’s Disease-Team Thomas Biederer ASAP 020616 through the Michael J Fox Foundation for Parkinson’s Research to LVD, and Alzheimer’s of Central Alabama.

## Author Contributions

**Nolwazi Z Gcwensa**: Conceptualization, Methodology, Formal analysis, Investigation, Data Curation, Writing – Original Draft; Writing – Review & Editing; Visualization. **Dreson L. Russell:** Methodology, Investigation, Data Curation, Writing – Original Draft. **Khaliah Y. Long:** Methodology, Investigation, Data Curation, Writing – Review & Editing. **Charlotte F. Brzozowski:** Methodology, Investigation, Writing – Original Draft, Writing – Review & Editing, Visualization. **Xinran Liu:** Conceptualization, Methodology, Investigation. **Karen L. Gamble:** Formal Analysis. **Rita M. Cowell:** Conceptualization, Writing – Review and Editing. **Laura A. Volpicelli-Daley:** Conceptualization, Methodology, Formal Analysis, Writing – Original Draft, Writing – Original Draft; Writing-Review & Editing, Visualization, Supervision, Funding Acquisition.

## Declaration of Generative AI and AI-assisted technologies in the Writing process

During the preparation of this work the author(s) used ChatGPT in order to help reword already established material and methods sections including: Animals, Intrastriatal injection of recombinant α-syn fibrils and Immunofluorescence and immunohistochemistry. After using this tool/service, the author(s) reviews and edited the content as need and take full responsibility for the content of the publication.

## References

Beesley, P. W., Herrera-Molina, R., Smalla, K. H., & Seidenbecher, C. (2014). The Neuroplastin adhesion molecules: key regulators of neuronal plasticity and synaptic function. J Neurochem, 131(3), 268–283. 10.1111/jnc.12816

Blumenstock, S., Rodrigues, E. F., Peters, F., Blazquez-Llorca, L., Schmidt, F., Giese, A., & Herms, J. (2017). Seeding and transgenic overexpression of alpha-synuclein triggers dendritic spine pathology in the neocortex. EMBO Mol Med, 9(5), 716–731. 10.15252/emmm.201607305

Bousset, L., Pieri, L., Ruiz-Arlandis, G., Gath, J., Jensen, P. H., Habenstein, B., Madiona, K., Olieric, V., Böckmann, A., Meier, B. H., & Melki, R. (2013). Structural and functional characterization of two alpha-synuclein strains. Nat Commun, 4, 2575. 10.1038/ncomms3575

Braak, H., Braak, E., Yilmazer, D., de Vos, R. A., Jansen, E. N., Bohl, J., & Jellinger, K. (1994). Amygdala pathology in Parkinson’s disease. Acta Neuropathol, 88(6), 493–500. 10.1007/bf00296485

Braak, H., & Del Tredici, K. (2017). Neuropathological Staging of Brain Pathology in Sporadic Parkinson’s disease: Separating the Wheat from the Chaff. J Parkinsons Dis, 7(s1), S71–s85. 10.3233/jpd-179001

Bridi, J. C., & Hirth, F. (2018). Mechanisms of α-Synuclein Induced Synaptopathy in Parkinson’s Disease. Front Neurosci, 12, 80. 10.3389/fnins.2018.00080

Carey, G., Görmezoğlu, M., de Jong, J. J. A., Hofman, P. A. M., Backes, W. H., Dujardin, K., & Leentjens, A. F. G. (2021). Neuroimaging of Anxiety in Parkinson’s Disease: A Systematic Review. Mov Disord, 36(2), 327–339. 10.1002/mds.28404

Chen, L., Nagaraja, C., Daniels, S., Fisk, Z. A., Dvorak, R., Meyerdirk, L., Steiner, J. A., Escobar Galvis, M. L., Henderson, M. X., Rousseaux, M. W. C., Brundin, P., & Chu, H. Y. (2022). Synaptic location is a determinant of the detrimental effects of α-synuclein pathology to glutamatergic transmission in the basolateral amygdala. Elife, 11. 10.7554/eLife.78055

Colom-Cadena, M., Pegueroles, J., Herrmann, A. G., Henstridge, C. M., Muñoz, L., Querol-Vilaseca, M., Martín-Paniello, C. S., Luque-Cabecerans, J., Clarimon, J., Belbin, O., Núñez-Llaves, R., Blesa, R., Smith, C., McKenzie, C. A., Frosch, M. P., Roe, A., Fortea, J., Andilla, J., Loza-Alvarez, P., … Lleó, A. (2017). Synaptic phosphorylated α-synuclein in dementia with Lewy bodies. Brain, 140(12), 3204–3214. 10.1093/brain/awx275

Earls, R. H., Menees, K. B., Chung, J., Barber, J., Gutekunst, C. A., Hazim, M. G., & Lee, J. K. (2019). Intrastriatal injection of preformed alpha-synuclein fibrils alters central and peripheral immune cell profiles in non-transgenic mice. J Neuroinflammation, 16(1), 250. 10.1186/s12974-019-1636-8

Froula, J. M., Castellana-Cruz, M., Anabtawi, N. M., Camino, J. D., Chen, S. W., Thrasher, D. R., Freire, J., Yazdi, A. A., Fleming, S., Dobson, C. M., Kumita, J. R., Cremades, N., & Volpicelli-Daley, L. A. (2019). Defining α-synuclein species responsible for Parkinson’s disease phenotypes in mice. J Biol Chem, 294(27), 10392–10406. 10.1074/jbc.RA119.007743

Froula, J. M., Henderson, B. W., Gonzalez, J. C., Vaden, J. H., McLean, J. W., Wu, Y., Banumurthy, G., Overstreet-Wadiche, L., Herskowitz, J. H., & Volpicelli-Daley, L. A. (2018). α-Synuclein fibril-induced paradoxical structural and functional defects in hippocampal neurons. Acta Neuropathol Commun, 6(1), 35. 10.1186/s40478-018-0537-x

Goralski, T. M., Meyerdirk, L., Breton, L., Brasseur, L., Kurgat, K., DeWeerd, D., Turner, L., Becker, K., Adams, M., Newhouse, D. J., & Henderson, M. X. (2024). Spatial transcriptomics reveals molecular dysfunction associated with cortical Lewy pathology. Nat Commun, 15(1), 2642. 10.1038/s41467-024-47027-8

Harding, A. J., Stimson, E., Henderson, J. M., & Halliday, G. M. (2002). Clinical correlates of selective pathology in the amygdala of patients with Parkinson’s disease. Brain, 125(Pt 11), 2431–2445. 10.1093/brain/awf251

Hornykiewicz, O. (1998). Biochemical aspects of Parkinson’s disease. Neurology, 51(2 Suppl 2), S2-9. 10.1212/wnl.51.2_suppl_2.s2

Huang, P., Xuan, M., Gu, Q., Yu, X., Xu, X., Luo, W., & Zhang, M. (2015). Abnormal amygdala function in Parkinson’s disease patients and its relationship to depression. J Affect Disord, 183, 263–268. 10.1016/j.jad.2015.05.029

Imbrosci, B., Schmitz, D., & Orlando, M. (2022). Automated Detection and Localization of Synaptic Vesicles in Electron Microscopy Images. eNeuro, 9(1). 10.1523/eneuro.0400-20.2021

Kilzheimer, A., Hentrich, T., Burkhardt, S., & Schulze-Hentrich, J. M. (2019). The Challenge and Opportunity to Diagnose Parkinson’s Disease in Midlife. Front Neurol, 10, 1328. 10.3389/fneur.2019.01328

Kordower, J. H., Olanow, C. W., Dodiya, H. B., Chu, Y., Beach, T. G., Adler, C. H., Halliday, G. M., & Bartus, R. T. (2013). Disease duration and the integrity of the nigrostriatal system in Parkinson’s disease. Brain, 136(Pt 8), 2419–2431. 10.1093/brain/awt192

Kouli, A., Torsney, K. M., & Kuan, W. L. (2018). Parkinson’s Disease: Etiology, Neuropathology, and Pathogenesis. In T. B. Stoker & J. C. Greenland (Eds.), Parkinson’s Disease: Pathogenesis and Clinical Aspects. Codon Publications Copyright: The Authors. 10.15586/codonpublications.parkinsonsdisease.2018.ch1

Kramer, M. L., & Schulz-Schaeffer, W. J. (2007). Presynaptic alpha-synuclein aggregates, not Lewy bodies, cause neurodegeneration in dementia with Lewy bodies. J Neurosci, 27(6), 1405–1410. 10.1523/jneurosci.4564-06.2007

Kumaresan, M., & Khan, S. (2021). Spectrum of Non-Motor Symptoms in Parkinson’s Disease. Cureus, 13(2), e13275. 10.7759/cureus.13275

Luk, K. C., Kehm, V., Carroll, J., Zhang, B., O’Brien, P., Trojanowski, J. Q., & Lee, V. M. (2012). Pathological α-synuclein transmission initiates Parkinson-like neurodegeneration in nontransgenic mice. Science, 338(6109), 949–953. 10.1126/science.1227157

Matuskey, D., Tinaz, S., Wilcox, K. C., Naganawa, M., Toyonaga, T., Dias, M., Henry, S., Pittman, B., Ropchan, J., Nabulsi, N., Suridjan, I., Comley, R. A., Huang, Y., Finnema, S. J., & Carson, R. E. (2020). Synaptic Changes in Parkinson Disease Assessed with in vivo Imaging. Ann Neurol, 87(3), 329–338. 10.1002/ana.25682

Opara, J. A., Brola, W., Leonardi, M., & Błaszczyk, B. (2012). Quality of life in Parkinson’s disease. J Med Life, 5(4), 375–381.

Parra-Rivas, L. A., Madhivanan, K., Aulston, B. D., Wang, L., Prakashchand, D. D., Boyer, N. P., Saia-Cereda, V. M., Branes-Guerrero, K., Pizzo, D. P., Bagchi, P., Sundar, V. S., Tang, Y., Das, U., Scott, D. A., Rangamani, P., Ogawa, Y., & Subhojit, R. (2023). Serine-129 phosphorylation of α-synuclein is an activity-dependent trigger for physiologic protein-protein interactions and synaptic function. Neuron, 111(24), 4006–4023.e4010. 10.1016/j.neuron.2023.11.020

Poewe, W. (2008). Non-motor symptoms in Parkinson’s disease. Eur J Neurol, 15 Suppl 1, 14–20. 10.1111/j.1468-1331.2008.02056.x

Qu, M., Gao, B., Jiang, Y., Li, Y., Pei, C., Xie, L., Zhang, Y., Song, Q., & Miao, Y. (2024). Atrophy patterns in hippocampus and amygdala subregions of depressed patients with Parkinson’s disease. Brain Imaging Behav. 10.1007/s11682-023-00844-9

Ramalingam, N., Jin, S. X., Moors, T. E., Fonseca-Ornelas, L., Shimanaka, K., Lei, S., Cam, H. P., Watson, A. H., Brontesi, L., Ding, L., Hacibaloglu, D. Y., Jiang, H., Choi, S. J., Kanter, E., Liu, L., Bartels, T., Nuber, S., Sulzer, D., Mosharov, E. V., … Dettmer, U. (2023). Dynamic physiological α-synuclein S129 phosphorylation is driven by neuronal activity. NPJ Parkinsons Dis, 9(1), 4. 10.1038/s41531-023-00444-w

Salzman, C. D., & Fusi, S. (2010). Emotion, cognition, and mental state representation in amygdala and prefrontal cortex. Annu Rev Neurosci, 33, 173–202. 10.1146/annurev.neuro.051508.135256

Šimić, G., Tkalčić, M., Vukić, V., Mulc, D., Španić, E., Šagud, M., Olucha-Bordonau, F. E., Vukšić, M., & P, R. H. (2021). Understanding Emotions: Origins and Roles of the Amygdala. Biomolecules, 11(6). 10.3390/biom11060823

Sorrentino, Z. A., Goodwin, M. S., Riffe, C. J., Dhillon, J. S., Xia, Y., Gorion, K. M., Vijayaraghavan, N., McFarland, K. N., Golbe, L. I., Yachnis, A. T., & Giasson, B. I. (2019). Unique α-synuclein pathology within the amygdala in Lewy body dementia: implications for disease initiation and progression. Acta Neuropathol Commun, 7(1), 142. 10.1186/s40478-019-0787-2

Stoyka, L. E., Arrant, A. E., Thrasher, D. R., Russell, D. L., Freire, J., Mahoney, C. L., Narayanan, A., Dib, A. G., Standaert, D. G., & Volpicelli-Daley, L. A. (2020). Behavioral defects associated with amygdala and cortical dysfunction in mice with seeded α-synuclein inclusions. Neurobiol Dis, 134, 104708. 10.1016/j.nbd.2019.104708

Taoufik, E., Kouroupi, G., Zygogianni, O., & Matsas, R. (2018). Synaptic dysfunction in neurodegenerative and neurodevelopmental diseases: an overview of induced pluripotent stem-cell-based disease models. Open Biol, 8(9). 10.1098/rsob.180138

Tye, K. M., Prakash, R., Kim, S. Y., Fenno, L. E., Grosenick, L., Zarabi, H., Thompson, K. R., Gradinaru, V., Ramakrishnan, C., & Deisseroth, K. (2011). Amygdala circuitry mediating reversible and bidirectional control of anxiety. Nature, 471(7338), 358–362. 10.1038/nature09820

Vargas, K. J., Makani, S., Davis, T., Westphal, C. H., Castillo, P. E., & Chandra, S. S. (2014). Synucleins regulate the kinetics of synaptic vesicle endocytosis. J Neurosci, 34(28), 9364–9376. 10.1523/jneurosci.4787-13.2014

Villalba, R. M., & Smith, Y. (2011). Differential structural plasticity of corticostriatal and thalamostriatal axo-spinous synapses in MPTP-treated Parkinsonian monkeys. J Comp Neurol, 519(5), 989–1005. 10.1002/cne.22563

Villar-Conde, S., Astillero-Lopez, V., Gonzalez-Rodriguez, M., Saiz-Sanchez, D., Martinez-Marcos, A., Ubeda-Banon, I., & Flores-Cuadrado, A. (2023). Synaptic Involvement of the Human Amygdala in Parkinson’s Disease. Mol Cell Proteomics, 22(12), 100673. 10.1016/j.mcpro.2023.100673

Volpicelli-Daley, L. A., Luk, K. C., & Lee, V. M. (2014). Addition of exogenous α-synuclein preformed fibrils to primary neuronal cultures to seed recruitment of endogenous α-synuclein to Lewy body and Lewy neurite-like aggregates. Nat Protoc, 9(9), 2135–2146. 10.1038/nprot.2014.143

Volpicelli-Daley, L. A., Luk, K. C., Patel, T. P., Tanik, S. A., Riddle, D. M., Stieber, A., Meaney, D. F., Trojanowski, J. Q., & Lee, V. M. (2011). Exogenous α-synuclein fibrils induce Lewy body pathology leading to synaptic dysfunction and neuron death. Neuron, 72(1), 57–71. 10.1016/j.neuron.2011.08.033

Wang, J., Sun, L., Chen, L., Sun, J., Xie, Y., Tian, D., Gao, L., Zhang, D., Xia, M., & Wu, T. (2023). Common and distinct roles of amygdala subregional functional connectivity in non-motor symptoms of Parkinson’s disease. NPJ Parkinsons Dis, 9(1), 28. 10.1038/s41531-023-00469-1

Yamazaki, M., Arai, Y., Baba, M., Iwatsubo, T., Mori, O., Katayama, Y., & Oyanagi, K. (2000). Alpha-synuclein inclusions in amygdala in the brains of patients with the parkinsonism-dementia complex of Guam. J Neuropathol Exp Neurol, 59(7), 585–591. 10.1093/jnen/59.7.585

Zhang, J., Chen, R., Shi, F., Yang, P., Sun, K., Yang, X., & Jin, Y. (2021). Genome-wide data mining to construct a competing endogenous RNA network and reveal the pivotal therapeutic targets of Parkinson’s disease. J Cell Mol Med, 25(13), 5912–5923. 10.1111/jcmm.16190

Zhang, L., Shao, Y., Tang, C., Liu, Z., Tang, D., Hu, C., Liang, X., Hu, Z., & Luo, G. (2022). Identification of Novel Biomarkers in Platelets for Diagnosing Parkinson’s Disease. Eur Neurol, 85(2), 122–131. 10.1159/000520102

Zhang, W. H., Zhang, J. Y., Holmes, A., & Pan, B. X. (2021). Amygdala Circuit Substrates for Stress Adaptation and Adversity. Biol Psychiatry, 89(9), 847–856. 10.1016/j.biopsych.2020.12.026

Zhao, N., Yang, Y., Zhang, L., Zhang, Q., Balbuena, L., Ungvari, G. S., Zang, Y. F., & Xiang, Y. T. (2021). Quality of life in Parkinson’s disease: A systematic review and meta-analysis of comparative studies. CNS Neurosci Ther, 27(3), 270–279. 10.1111/cns.13549

