## Supplementary Information for "Cortico-amygdala synaptic structural abnormalities produced by templated aggregation of α-synuclein"

### *Supplemental Information*

**Whole-brain images of MON and PBS-injected Mice 6- and 12-weeks post-injections**

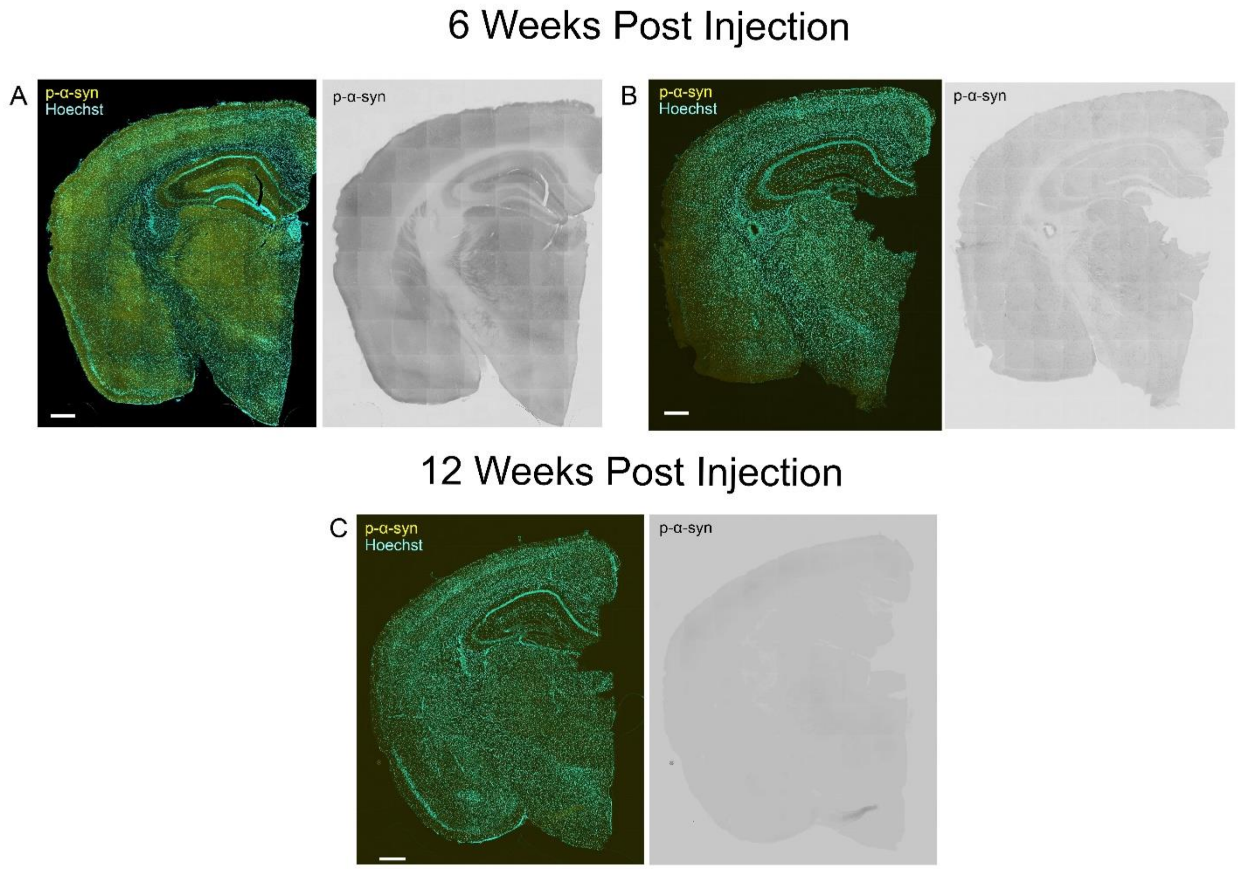


**Supplemental Figure 1.** MON and PBS injection does not induce α-synuclein inclusions in mouse BLA 6- and 12-weeks post-injection. Representative images for 3 – 4 month old mice injected with (A, C) phosphate buffered saline (PBS) or (B) monomeric α-synuclein (MON) and sacrificed 6- and 12- weeks post-injection. Left panels show p-Ser129-α-synuclein stain in (yellow) and Hoechst stained nuclei (blue). No p-α-synuclein inclusions are observed. Scale Bar = 500 μm.

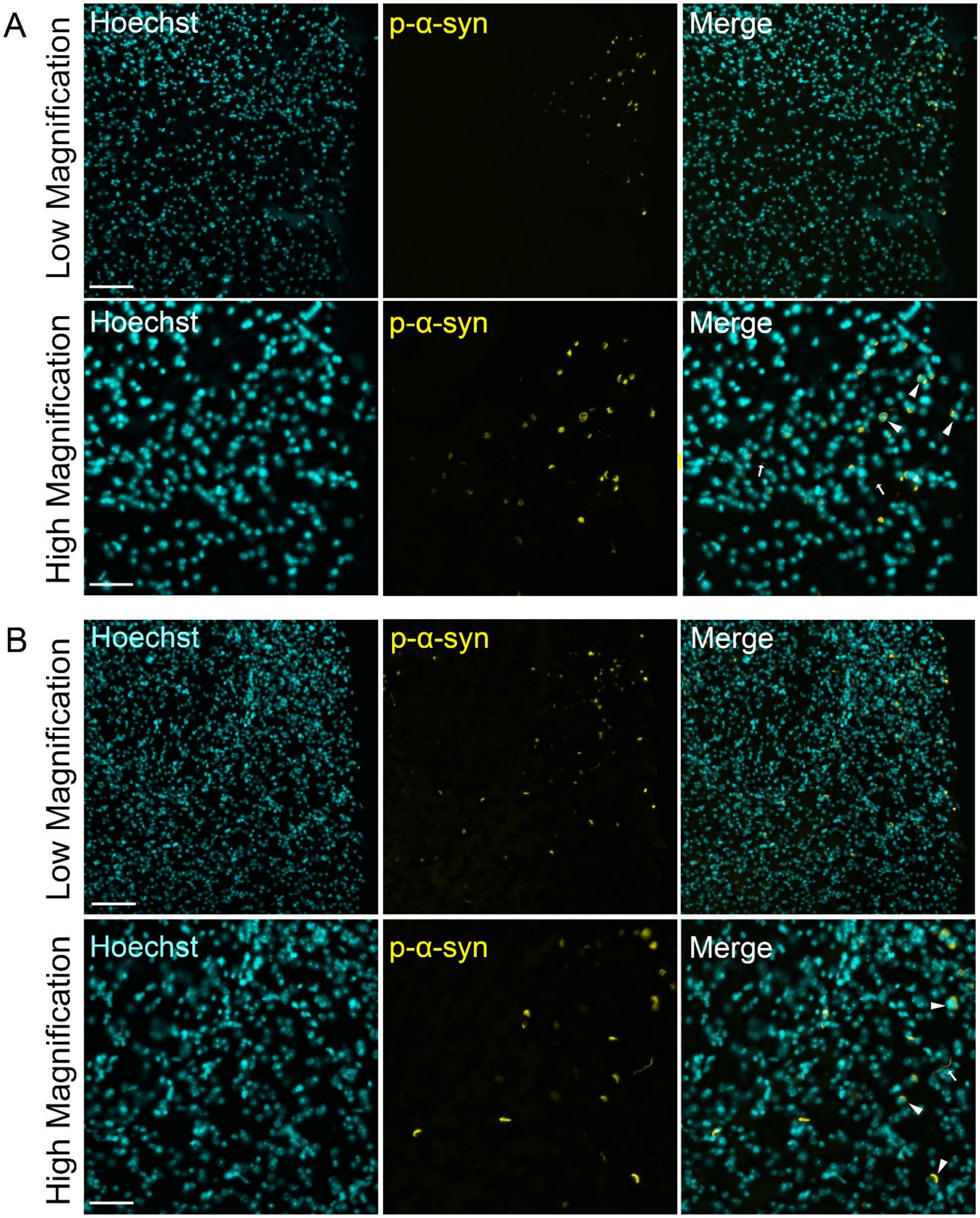
A
Low Magnification
High Magnification
Hoechst
p-α-syn
Merge
Hoechst
p-α-syn
Merge
B
Low Magnification
High Magnification
Hoechst
p-α-syn
Merge
Hoechst
p-α-syn
Merge

**Supplemental Figure 2.** Low (20X) and high (40X) magnification images of thalamic p-α-synuclein inclusions in PFF-injected mice (A) 6-weeks post-injection and (B) 12-weeks post-injection with arrows to show pSer127-α-syn+ (yellow) intracellular inclusions overlapping with Hoechst stained nuclei (cyan). Lewy body-like inclusions indicated by white arrowheads and Lewy neurite-like inclusions indicated by white arrows. Scale Bar = 100 μm and 50 μm for low magnification and high magnification, respectively.

**Mean densities and mean volumes of total of VGLUT1 and HOMER1 puncta at 6 weeks post-injection**

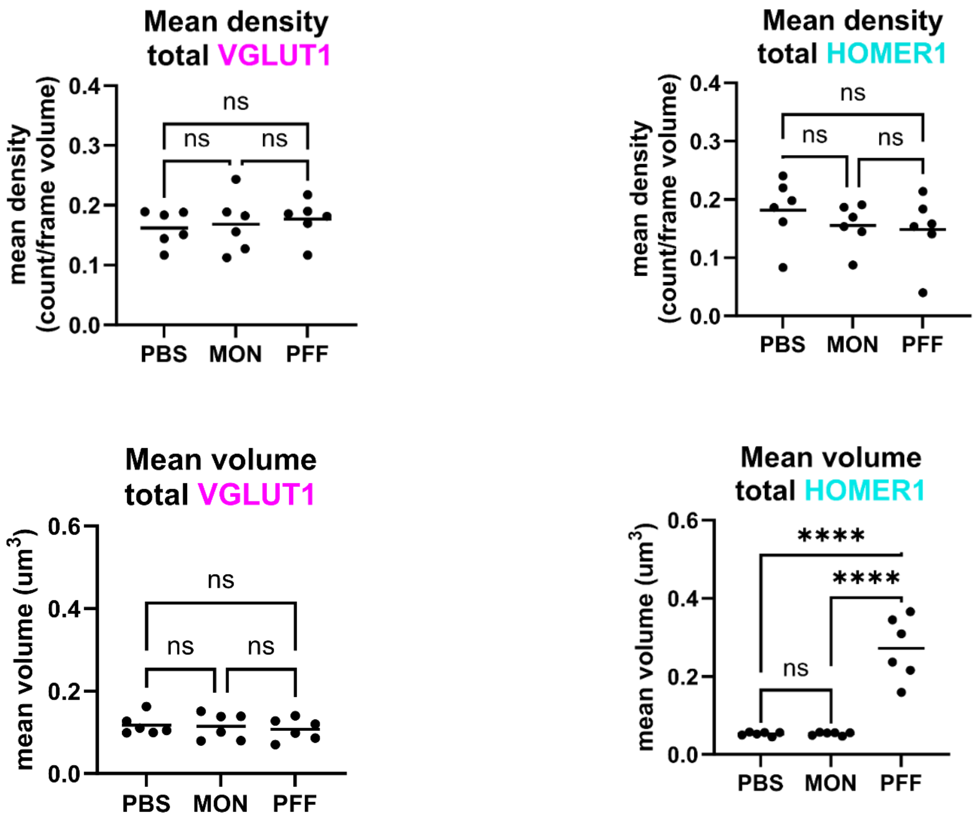


**Supplemental Figure 3.** Effect of p-a-synuclein inclusion formation in density and volume of cortico-amygdala total surfaces at 6 weeks post-injection (A) Mice were injected with either PBS, monomeric α –synuclein (MON), or PFFs and sacrificed 6 weeks post-injection. Mean values of the puncta density for (upper panels) total VGLUT1+ and total HOMER1+ puncta showed overall no significant differences. Mean values for volume of (lower panels) total VGLUT1+ puncta showed no significant differences between groups. Total volume of HOMER1+ puncta in PFF-injected animals was significantly larger than control. Statistical model: One-way ANOVA with Brown-Forsythe correction applied to groups with unequal variance ($p > 0.05$). Data points represent average values for individual mouse

**CORTICO-AMYGDALA PROJECTIONS**

Table S1A. Statistics summary table for mean density of total VGLUT1+ and total HOMER1+ puncta 6 weeks post-injection

| | ***Test*** | ***[group] (mean, SD)*** | ***F**** | ***Dfn*** | ***Dfd*** | ***p*** |
|---|---|---|---|---|---|---|
| **TOTAL VGLUT1** | One-way ANOVA | PBS (0.1622, 0.0296)<br>MON (0.1685, 0.0473)<br>PFF (0.1770, 0.0334) | 0.2343 | 2 | 15 | 0.7939 |
| **TOTAL HOMER1** | One-way ANOVA | PBS (0.1816, 0.0553)<br>MON (0.1554, 0.0379)<br>PFF (0.1484, 0.0590) | 0.6918 | 2 | 15 | 0.5159 |

Table S1B. Statistics summary table for mean volume of total VGLUT1+ and total HOMER1+ puncta 6 weeks post-injection

| | ***Test*** | ***[group] (mean, SD)*** | ***F**** | ***Dfn*** | ***Dfd*** | ***p*** |
|---|---|---|---|---|---|---|
| **TOTAL VGLUT1** | One-way ANOVA | PBS (0.1170, 0.0245)<br>MON (0.1146, 0.0320)<br>PFF (0.1070, 0.0266) | 0.2098 | 2 | 15 | 0.8131 |
| **TOTAL HOMER1** | Brown-Forsyth and One-way Welch’s ANOVA | PBS (0.0527, 0.0049)<br>MON (0.0531, 0.0042)<br>PFF (0.2720, 0.0805) | 44.15 | 2 | 5.064 | 0.0006 |

**Mean densities and mean volumes of total of VGLUT1 and HOMER1 puncta at 12 weeks post-injection**

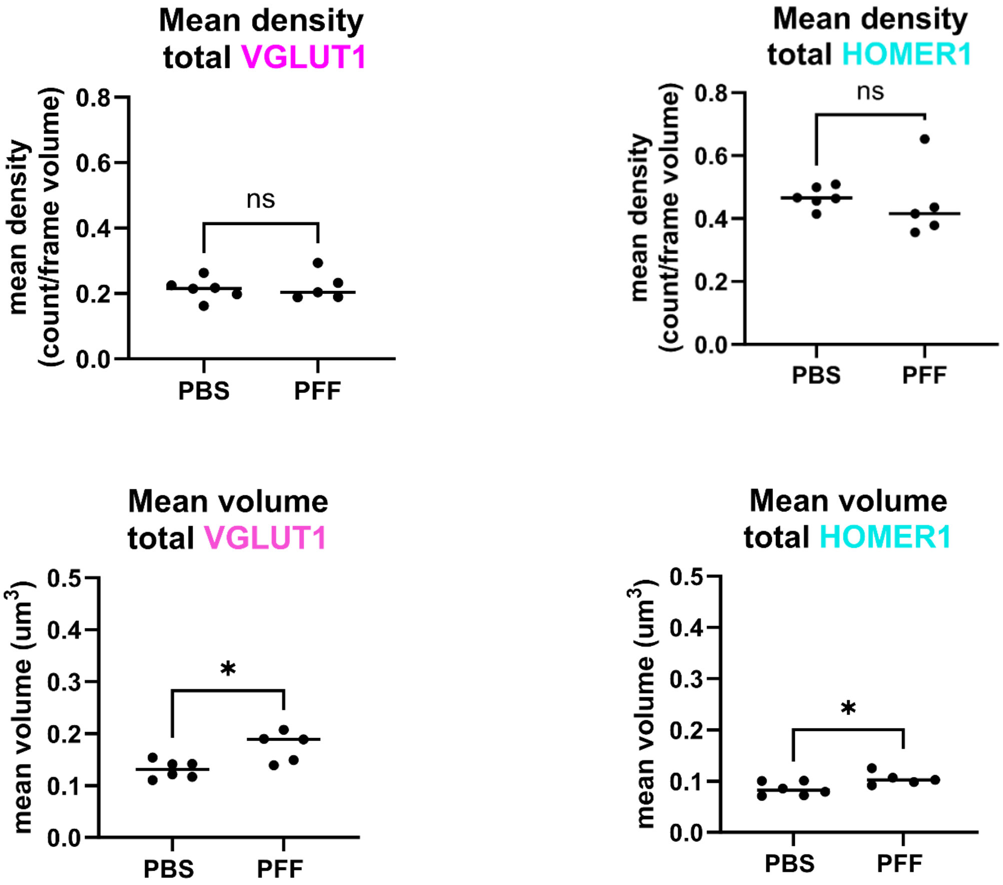


**Supplemental Figure 4.** Effect of p-a-synuclein inclusion formation in density and volume of cortico-amygdala total surfaces at 12 weeks post-injection. (A) Mice were injected with either PBS, monomeric α –synuclein (MON), or PFFs and sacrificed 6 weeks post-injection. Mean values of the puncta density (upper panels) for total VGLUT1+ and total HOMER1+ puncta showed overall no significant differences. Mean values for volume of (lower panels) total VGLUT1+ puncta and total HOMER1+ puncta showed significant increases in PFF-injected animals was significantly larger than control. Statistical model: Students t-test with Welch's correction applied to groups with significant differences in variance (p > 0.5). Data points represent average values for individual mouse.

| **CORTICO-AMYGDALA PROJECTIONS** | | | | | |
|---|---|---|---|---|---|
| Table S2A. Statistics summary table for mean density of total VGLUT1+ and total HOMER1+ puncta 12 weeks post-injection | | | | | |
| | ***Test*** | ***[group] (mean, SD)*** | ***t*** | ***df*** | ***p (p < 0.05)*** |
| **TOTAL VGLUT1** | Parametric Student's t-test | PBS (0.2134, 0.0333) PFF (0.2215, 0.0443) | 0.3493 | 9 | 0.7349 |
| **TOTAL HOMER1** | Parametric Student's t-test, Welch's correction | PBS (0.4684, 0.0337) PFF (0.4480, 0.1188) | 0.3720 | 4.536 | 0.7266 |
| Table S2B. Statistics summary table for mean volume of total VGLUT1+ and total HOMER1+ puncta 12 weeks post-injection | | | | | |
| | ***Test*** | ***[group] (mean, SD)*** | ***t*** | ***df*** | ***p (p < 0.05)*** |
| **TOTAL VGLUT1** | Parametric Student's t-Test | PBS (0.1309, 0.0171) PFF (0.1747, 0.0292) | 3.110 | 9 | **0.0125** |
| **TOTAL HOMER1** | Parametric Student's t-Test | PBS (0.0849, 0.0132) PFF (0.1049, 0.0127) | 2.552 | 9 | **0.0311** |

**Mean densities and mean volume of total VGLUT2 and HOMER1 PUNCTA at 6 weeks post-injection**

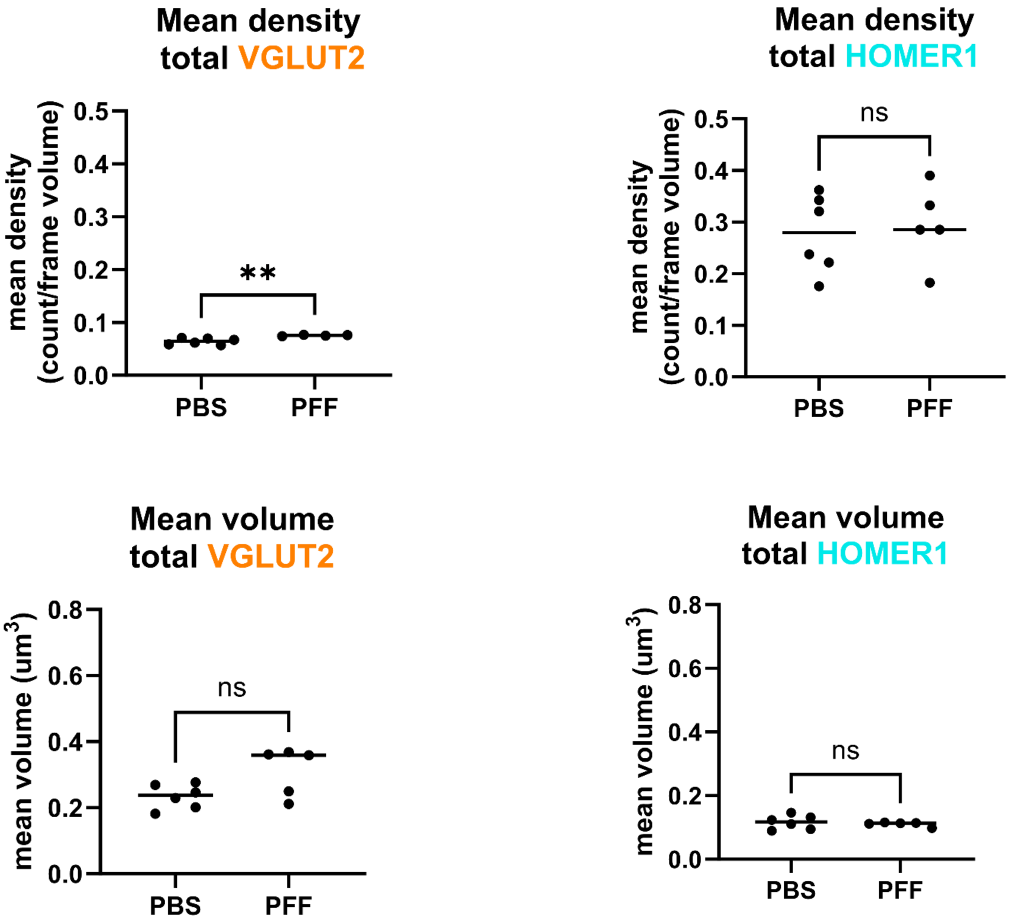


**Supplemental Figure 5.** Effect of p-a-synuclein inclusion formation in density and volume of thalamo-amygdala total surfaces at 6 weeks post-injection. (A) Mice were injected with either PBS, monomeric α –synuclein (MON), or PFFs and sacrificed 6 weeks post-injection. Mean values of the puncta density (upper panels) for total VGLUT2+ and HOMER1+ puncta. Analysis showed significant increase in density of VGLUT2+ terminals and showed no significant differences for total HOMER1+ puncta. Mean values for volume of (lower panels) total VGLUT1+ puncta and total HOMER1+ puncta showed no significant differences between groups. Statistical model: Students t-test with Welch's correction applied to groups with significant differences in variance ($p > 0.5$). Data points represent average values for individual mouse.

**THALAMO-AMYGDALA PROJECTIONS**

Table S3A. Statistics summary table for mean density of total VGLUT2+ and total HOMER1+ puncta 6 weeks post-injection

| | ***Test*** | ***[group] (mean, SD)*** | ***t*** | ***df*** | ***p (p < 0.05)*** |
|---|---|---|---|---|---|
| **TOTAL VGLUT2** | Parametric Student’s t-Test, Welch’s correction | PBS (0.0641, 0.0058)<br>PFF (0.0753, 0.0012) | 4.581 | 5.600 | **0.0045** |
| **TOTAL HOMER1** | Parametric Student’s t-Test | PBS (0.2767, 0.0753)<br>PFF (0.2950, 0.0762) | 0.3984 | 9 | 0.6996 |

Table S3B. Statistics summary table for mean volume of total VGLUT2+ and total HOMER1+ puncta 6 weeks post-injection

| | ***Test*** | ***[group] (mean, SD)*** | ***t*** | ***df*** | ***p (p < 0.05)*** |
|---|---|---|---|---|---|
| **TOTAL VGLUT2** | Parametric Student’s t-Test | PBS (0.2340, 0.0374)<br>PFF (0.3100, 0.0738) | 2.221 | 9 | 0.0535 |
| **TOTAL HOMER1** | Parametric Student’s t-Test | PBS (0.1162, 0.0220)<br>PFF (0.1104, 0.0073) | 0.5596 | 9 | 0.5894 |

**Mean densities and mean volume of total VGLUT2 and HOMER1 PUNCTA at 12 weeks post-injection**

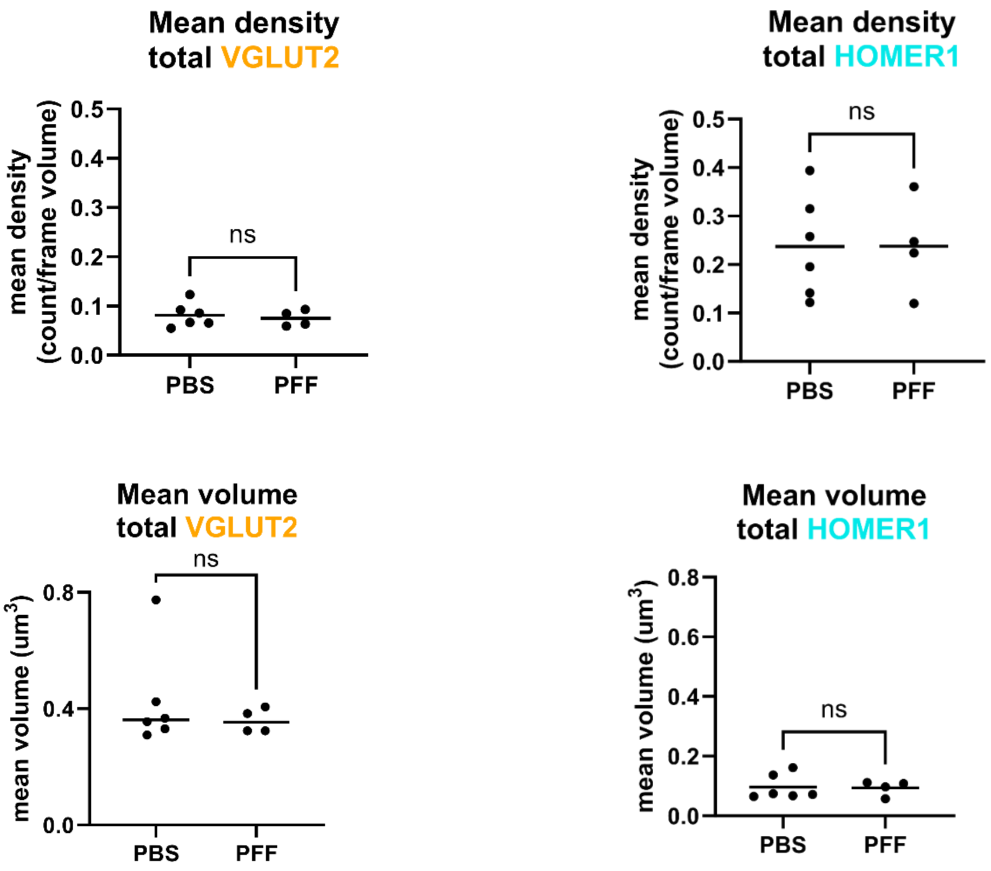


**Supplemental Figure 6.** Effect of p-a-synuclein inclusion formation in density and volume of thalamo-amygdala total surfaces at 12 weeks post-injection. (A) Mice were injected with either PBS, monomeric α –synuclein (MON), or PFFs and sacrificed 6 weeks post-injection. Mean values of the puncta density (upper panels) for total VGLUT2+ and HOMER1+ puncta. Analysis showed no significant difference in density of VGLUT2+ terminals and showed no significant differences for total HOMER1+ puncta. Mean values for volume of (lower panels) total VGLUT1+ puncta and total HOMER1+ puncta showed no significant differences between groups. Statistical model: Students t-test with Welch's correction applied to groups with significant differences in variance ($p > 0.5$). Data points represent average values for individual mouse.

**THALAMO-AMYGDALA PROJECTIONS**

Table S4A. Statistics summary table for mean density of total VGLUT2+ and total HOMER1+ puncta 12 weeks post-injection

| | ***Test*** | ***[group] (mean, SD)*** | ***t*** | ***df*** | ***p (p < 0.05)*** |
|---|---|---|---|---|---|
| **TOTAL VGLUT2** | Parametric Student’s t-test | PBS (0.0812, 0.0248) PFF (0.0749, 0.0162) | 0.4429 | 8 | 0.6696 |
| **TOTAL HOMER1** | Parametric Student’s t-test | PBS (0.2373, 0.1052) PFF (0.2377, 0.0989) | 0.007152 | 8 | 0.9945 |

Table S4B. Statistics summary table for mean volume of total VGLUT2+ and total HOMER1+ puncta 12 weeks post-injection

| | ***Test*** | ***[group] (mean, SD)*** | ***t*** | ***df*** | ***p (p < 0.05)*** |
|---|---|---|---|---|---|
| **TOTAL VGLUT2** | Parametric Student’s t-test | PBS (0.3575, 0.0432) PFF (0.3595 0.0421) | 0.06808 | 7 | 0.9476 |
| **TOTAL HOMER1** | Parametric Student’s t-test | PBS (0.0961, 0.0420) PFF (0.0934, 0.0250) | 0.1147 | 8 | 0.9115 |

**Distributions of synaptic vesicle area and synaptic bouton lengths**

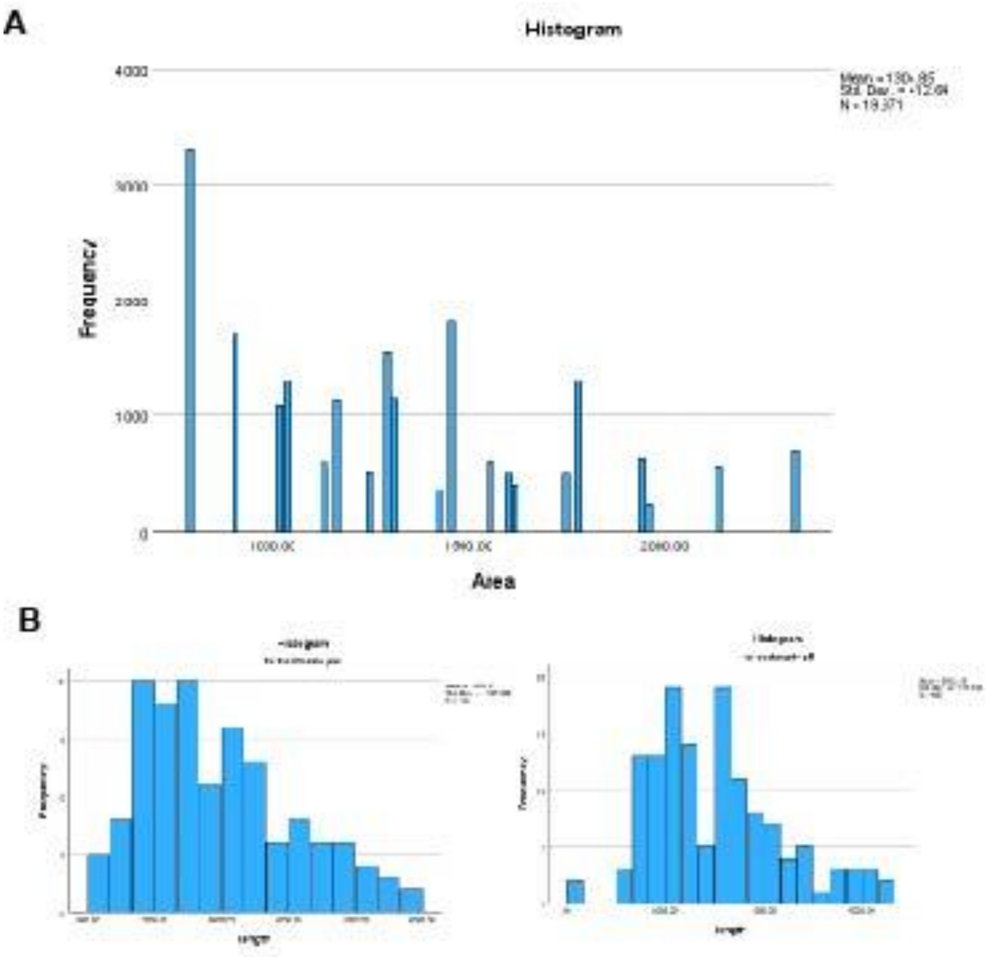


**Supplemental Figure 7. (A)** Distributions of synaptic vesicle areas shows a non-continuous distribution because of the thresholding built-in to the neural network. This distribution prompted the authors to bin the synaptic vesicle area data into areas above and below the grand median. (B). Distributions of synaptic bouton lengths for PBS and PFF treated mice. The PFF treated mice show a bimodal distribution suggesting that there may be larger synaptic boutons, but the linear mixed model analyses were not significant.
